# Whole-Brain High-Resolution Metabolite Mapping with 3D Compressed-Sensing-SENSE-LowRank ^1^H FID-MRSI

**DOI:** 10.1101/2020.05.18.101618

**Authors:** Antoine Klauser, Paul Klauser, Frédéric Grouiller, Sebastien Courvoisier, Francois Lazeyras

## Abstract

There is a growing interest in the neuroscience community to map the distribution of brain metabolites in vivo. Magnetic resonance spectroscopic imaging (MRSI) is often limited by either a poor spatial resolution and/or a long acquisition time which severely limits its applications for clinical or research purposes. Building on a recently developed technique of acquisition-reconstruction for 2D MRSI, we combined fast Cartesian ^1^H-FID-MRSI acquisition sequence, compressed-sensing acceleration, and low-rank total-generalized-variation constrained reconstruction to produce 3D high-resolution whole-brain MRSI with a significant acquisition time reduction. We first evaluated the acceleration performance using retrospective undersampling of a fully-sampled dataset. Second, a 20 min accelerated MRSI acquisition was performed on the brain of three healthy volunteers resulting in metabolite maps with 5 mm isotropic resolution. The metabolite maps exhibited the detailed neurochemical composition of all brain regions and revealed parts of the underlying brain anatomy. The latter assessment used previous reported knowledge and a brain atlas-based analysis to show consistency of the concentration contrasts and ratio across all brain regions. These results acquired on a clinical 3 Tesla MRI successful combinae of the 3D ^1^H-FID-MRSI with a constrained reconstruction to produce detailed mapping of metabolite concentrations at high-resolution over the whole brain, with an acquisition time suitable for clinical or research settings.

## I. INTRODUCTION

Measuring metabolite distributions in three dimensions over the whole human brain using proton magnetic resonance spectroscopic imaging (^1^H-MRSI) has been the subject of two decades of intense research. From early multi-slice methods [1, 2] covering a large portion of the brain to the first development of spatio-spectral encoding (SSE) techniques [3–5] proposed by Mansfield [6], whole brain ^1^H-MRSI unravels distributions with unique patterns for each metabolite and provides original physiological information that complements the usual MR imaging. Since then, 3D ^1^H-MRSI methods for human brain imaging have been improved by implementing acceleration techniques [7–9] or significant increase in spatial resolution [10, 11]. More complete overview of ^1^H-MRSI techniques can be found in review articles [12–14].

Recently, advances in reconstruction methods have provided new solutions to the inherent limitations of ^1^H-MRSI. The ^1^H-MRSI acquisition geometry is often restricted to a rectangular volume-of-interest (VOI) with saturation of the outer signal to prevent skull lipid contamination. However,recent work on efficient post-acquisition lipid decontamination [15–18] enables the measurement of whole brain slices without limited VOI by in plane selective excitation. In addition, employing models with constraints of prior information for ^1^H-MRSI data reconstruction significantly improves spectral quality and the resulting metabolite distributions [19, 20]. A priori knowledge of the underlying signal can also be exploited for super-resolution [21, 22] or drastic acceleration of ^1^H-MRSI acquisition [23, 24]. Acquisition of whole-brain MRSI using traditional methods could be particularly lengthy due to the necessity to encode the large 4D k-t space. However, acquisition time can be significantly reduced by using several techniques such as the free induction decay (FID)-MRSI acquisition, parallel imaging, and compressed sensing (CS). The FID-MRSI sequence first known for its application in ^31^P and other nuclei spectroscopic imaging [25, 26], has been more recently proposed for ^1^H-MRSI [27, 28]. The simple sequence design dramatically shortens acquisition time compared with usual spin-echo methods by allowing low-flip-angle excitation and sub-second repetition time (TR). Implementing of parallel imaging techniques in 3D ^1^H-MRSI of the human brain can reduce the acquisition time by a factor of 2 to 8. [8, 9, 29]. Following the recent developments of CS for MRI [30, 31], the sparsity present in the 3D MRSI signal can also be exploited to accelerate the encoding of the 4D k-t space. The effectiveness of this approach was demonstrated for ^13^C or ^19^F [32, 33] as well as in combination with fast spatial-spectral encoding [34]. Applying CS to ^1^H-MRSI of the human brain grants accelerations up to a factor of 4 but has been limited to 2D acquisitions so far [17, 35–38].

State-of-the-art techniques offering 3D whole-brain ^1^H-MRSI often rely on the use of SSE [14] which uses entire lines to encode the 4D k-t space. Hence, these approaches allow highly accelerated acquisitions and permit MRSI acquisitions of the whole-brain in high-resolution within a duration compatible with clinical requirements. The trajectory can be either echo-planar [24, 39], concentric circles [10, 40], rosette [41] or spiral [42]. With a signal-to-noise ratio (SNR) / time comparable to Cartesian encoding of same resolution [14], SSE technique might nevertheless result in low SNR data that requires averaging or specific denoising model-based approaches[24]. The method of acceleration presented hereafter relies on undersampling of the Cartesian 3D k-space. The non-linear reconstruction process [17] is expected to preserve SNR or to results in a slower decrease as function of the acceleration factor. On the other hand, high acceleration is expected to affect the spatial distributions of the MRSI signal as pointed out in previous studies of CS with MRI [43, 44]. In this study, we aim to evaluate the capability of the proposed framework as an alternative to the acceleration of 3D MRSI with SSE.

We present original results of high-resolution 3D ^1^H-MRSI of the whole human brain on a 3T clinical system. The acquisition-reconstruction scheme combines a fast FID-MRSI high-resolution sequence accelerated by random undersampling and reconstructed with a CS SENSE low-rank model (CS-SENSE-LR [17]) extended to 3D. This comprehensive approach, combining FID and parallel acquisition schemes to CS-SENSE-LR reconstruction aims to drastically shorten the acquisition time and allows the implementation of high-resolution 3D MRSI into clinical and research scanning protocols. High-resolution metabolite mapping will have a great range of applications from neurology to psychiatry [45, 46]. In clinical neuroscience and with usual spectroscopic techniques, interpretation of spectroscopy findings is limited by the VOI approach, neglecting large cerebral parts around the volume selected by the a priori hypothesis. There is a strong need for whole-brain spectroscopic imaging which allows not only the mapping of metabolites in all brain regions but also the integration of brain spectroscopic data with other imaging modalities including structural and diffusion MR [47].

## II. METHODS

### A. Sequence and Acquisition

A ^1^H-FID-MRSI [27, 28] sequence with a 3 dimensional phase encoding was implemented (Fig. 1) on a 3T Prisma fit MRI (Siemens, Erlangen, Germany) using a receiver head coil with *N*_*c*_ = 64 elements. A Shinnar-LeRoux optimized slab-selective excitation pulse of 0.9 ms was used with a 9.5-kHz bandwidth. Water suppression enhance through *T*_1_ effects (WET) [48] [48] was implemented using three Gaussian pulses with 60 Hz bandwidth and separated by 20 ms each. The acquisition delay between the excitation and the signal acquisition was minimized to an echo time (TE) of 0.65 ms. A 4-kHz sampling rate FID was acquired with *T* = 1024 points followed by spoiler gradients for a TR of 355 ms. To avoid signal saturation considering the maximum *T*_1_ value among metabolite to be 1400 ms [49], the excitation flip angle was set to 35 degrees. To avoid aliasing from signal outside the slab, two 30-mm-thick outer-volume saturation bands (OVS) were positioned directly below and above the slab. The excited slab size was (A/P-R/L-H/F) 210 mm by 160 mm by 95 mm. The 3D encoding volume was set slightly larger to 210 mm by 160 mm by 105 mm again to prevent aliasing. An encoding matrix of 42 × 32 × 20 resulted in a 131*µl* voxel volume (5 mm isotropic). A fast reference water measurement was performed to determine the coil sensitivity profiles, with acquisition parameters identical to the main FID-MRSI sequence but without WET water suppression and with lower resolution 32×24×16 (6.6 mm isotropic). The same encoding volume and slab were used but with 31-ms TR, 48 points in FID and a 5-degrees excitation flip angle. An anatomical 3D *T*_1_-weighted MPRAGE sequence was also acquired during the session for navigation purpose and for anatomical segmentation of the volunteers’ brain.

**FIG. 1.**
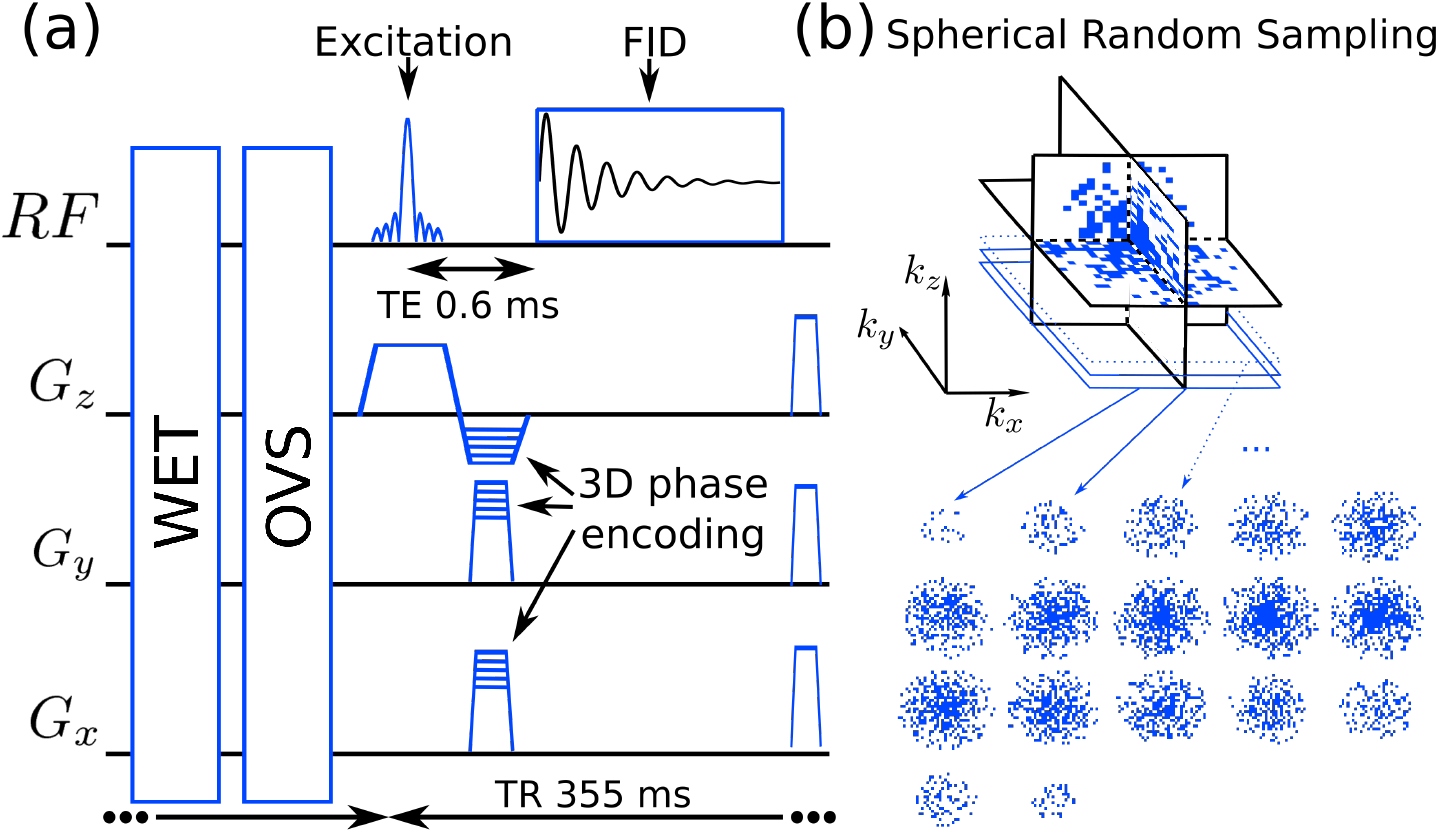
Schematic of the FID-MRSI sequence with 3D phase encoding (left) preceded by the water suppression enhanced through T1 effects (WET) and outer-volume saturation bands (OVS) sequence blocks. An example of 3D undersampled k-space by factor 3.5 is shown (right)

#### Sparse and random 3D phase-encoding

MRSI raw data were acquired sparsely and randomly over the three-dimensional Fourier domain (k-space) to enable CS acceleration. The sequential phase encoding method of the FID-MRSI sequence allows for a straight-forward implementation of the random k-space sampling. During sequence preparation, a 3D mask was computed containing only k-space coordinates to be acquired. Defining 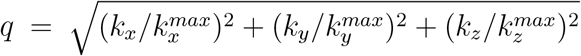, the random sampling of the 3D mask was constrained to a density distribution following *q*^−1^ but with a fully-sampled spheroid of radius 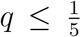 (Fig. 1). During the sequence acquisition, following the preparation, FID MRSI data were acquired according to the 3D k-space mask.

### B. MRSI data preprocessing and reconstruction model

#### Remaining water and lipid signal removal

Residual water was removed using Hankel singular value decomposition (HSVD) method [50]. HSVD was applied separately to each coil element and time series in the acquired points of the k-space. The lipid suppression was performed in k-space on the signal from each coil element using a lipid subspace projection method [17] based on the metabolite-lipid spectral orthogonality hypothesis [15]. The computation time of the residual water removal was below 60 min while the lipid suppression was computed within 2 min using Matlab R2018b (The MathWorks, Inc., Natick, Massachusetts, US) and 16 cores 3.00GHz Intel(R)Xeon(R)-E5 CPU. The methods are described in detail in previously published work [17].

#### CS-SENSE-LR reconstruction

The CS approach requires reconstruction of the sparsely-sampled MRSI data with an appropriate model that maximizes data fidelity while constraining spatial sparsity in metabolite distributions [30, 31]. We employed here a low-rank constrained model that includes total generalized variation (TGV) regularization [51]. We describe hereafter briefly the 3D extension of the CS-SENSE-LR reconstruction that was previously described in [17]. MRSI raw data measured by phased array coil element *c* = 1, …, *N*_*c*_ at time *t* and at k-space coordinate **k** can be expressed with the forward model

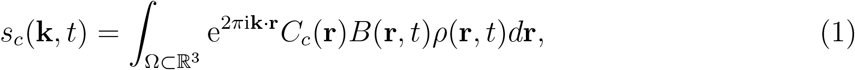

with 3D spatial coordinates **r** integrated over Ω the spatial support of the measured object and *ρ*(**r**, *t*) ∈ ℂ the transverse magnetization. *C*_*c*_(**r**) ∈ ℂ represent the coil sensitivity profiles and 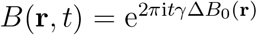 the spatial frequency shift caused by *γ*Δ*B*_0_(**r**), the magnetic field inhomogeneity map (in Hz). The goal of the reconstruction is to retrieve the original *ρ*(**r**, *t*) from the sparsely sampled signal *s*_*c*_(**k**, *t*) knowing *C*_*c*_(**r**) and Δ*B*_0_(**r**).

The low-rank assumption implies a decomposition of the transverse magnetization in *K* spatial and temporal components, *U*_*n*_(**r**), *V*_*n*_(*t*), *n* = 1, …, *K*:

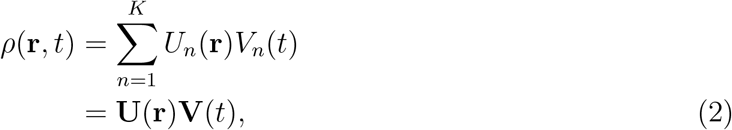

with the last line employing vectorial notation. Considering discrete spatial and temporal sampling points (*N*_*t*_ times points, *N*_*k*_ k-space acquisition points, *N*_*x*_, *N*_*y*_, *N*_*z*_ the reconstruction image size), *s*_*c*_(**k**, *t*), **U**(**r**)**V**(*t*) become multi-dimensional arrays **s, U, V** with size respectively (*N*_*c*_ × *N*_*k*_ × *N*_*t*_), (*N*_*x*_ × *N*_*y*_ × *N*_*z*_ × *K*) and (*K* × *N*_*t*_). The forward model (1) reads then

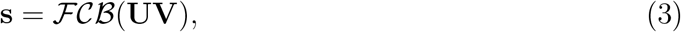

with the discrete Fourier transform operator ℱ and 𝒞 and ℬ operators applying *C*_*c*_(**r**) and *B*(**r**, *t*) on discrete coordinates. First, remaining water and lipid signal are removed from **s** as described above in previous section. Second, the magnetization in image space *ρ* = **UV**, is reconstructed from **s** with a 3D low-rank TGV model in [52]. The TGV minimization problem [51] enables the determination of the spatial and temporal components

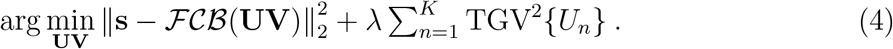

The coil sensitivity profiles, *C*_*c*_(**r**) were determined from the fast reference water acquisition using ESPIRiT [53] and Δ*B*_0_(**r**) was estimated using *multiple signal classification algorithm* (MUSIC) [54] on the water signal from the same fast reference scan. By including coil sensitivity profiles and the TGV regularization that imposes sparsity in spatial first and second-order gradients, the CS-SENSE-LR reconstruction permits a faithful reconstruction of randomly sampled MRSI data [43, 55]. As a consequence of the sequence design, there is a delay between the excitation and the FID acquisition that causes a first order phase in the reconstructed spectra. To correct this effect, the first 2 missing points of the FID were estimated using a reverse autoregressive model [56] in each voxel of the reconstructed MRSI dataset *ρ* = **UV** individually. The MRSI reconstruction including all pipeline steps takes approximately 12 hours to complete using Matlab R2018b (The MathWorks, Inc., Natick, Massachusetts, US) using a workstation with 16-cores 3.00GHz Intel(R)Xeon(R)-E5 CPU.

#### LCModel Quantification

In a last step, the reconstructed MRSI signal was fitted with LCModel [57, 58] to quantify metabolites voxel by voxel over the range of 1.0 to 4.2ppm. The LCModel basis was simulated using GAMMA package [59] and with parameters matching the acquisition sequence. The metabolites in the basis included: N-acetylaspartate, N-acetyl aspartylglutamate, creatine, phosphocreatine, glycerophospho-choline, phosphocholine, myo-inositol, scyllo-inositol, glutamate, glutamine, lactate, gamma-aminobutyric acid, glutathione, taurine, aspartate and alanine. LCModel results were expressed in institutional units (I.U.) that allow for comparison across metabolites and subjects (see LCModel documentation [58]). Although an absolute quantification of the metabolite signal would be possible with the proposed acquisition-reconstruction scheme, it would required multiple additional correction and adjustment factors such as tissue specific *T*_1_ corrections, water content correction and transmit amplitude for body and phased-array coils [60, 61]. It is however possible to give an estimate of the scaling between the I.U. and the metabolite concentrations in millimolar (mM). If we assume that *T*_1_ of the metabolites and the water is 1200 ms, and that the water concentration is 42.98mM in the whole brain [61], we can approximate that 1 I.U. ≈ 3.2 mM for all the data reported in this paper.

Lipid suppression by orthogonality as performed in this study might distort the baseline at 2ppm below the NAA singlet peak. To optimize LCModel fitting in presence of this distortion a singlet broad and inverse peak was added to the basis. It was simulated at 2 ppm with 20 Hz width and a *e*^*iπ*^ phase (opposite to NAA singlet). LCModel quantification provides also estimates of spectrum SNR and a Cramer-Rao Lower bound (CRLB) for each metabolite. These are usually considered as spectral quality metrics, but for the present results reconstructed with the CS-SENSE-LR model prior to the LCModel fitting, these parameters might not assess data quality properly. Especially as the accurate estimation of these parameters relies on the presence of measurement noise within the spectra. In the present work, the measurement noise is removed by using the denoising model. Therefore, CRLB values might be artificially low and SNR particularly high. Nevertheless, as they still reflect the fitting quality, they are provided in supplementary material. Instead, the average of the LCModel fit residuals is reported to describe spectral quality.

### C. Experiments

#### Assessing CS acceleration

To demonstrate the CS acceleration capability and to assess the metabolite maps reconstruction accuracy, a single MRSI acquisition without k-space undersampling was performed on a volunteer using the sequence details in II A with a full elliptical k-space encoding resulting in a 70 min acquisition. The raw data were first undersampled retrospectively to various extents to reproduce acceleration factors of 2, 3, 4, 5 or 6 and following the same variable density undersampling as the actual accelerated acquisition (described in II A). After undersampling of the raw data, metabolite maps were reconstructed following all the steps from II B. The TGV regularization parameter *λ* in (4) was adjusted on the fully-sampled dataset (3 × 10^−4^) and was kept the same for all acceleration factors. The reconstructed spectral quality was assessed at different accelerations and the fitting residuals of the LCModel were quantified as quality parameter. The resulting metabolite maps were compared qualitatively and quantitatively with a normalized root mean square error (RMSE) and structural similarity index (SSIM).

#### Healthy volunteers accelerated acquisition

The FID-MRSI with accelerated acquisition was performed and reconstructed with the CS-SENSE-LR model on three healthy volunteers to demonstrate reproducibility and robustness. Written informed consent was given by all the volunteers before participation and the study protocol was approved by the institutional ethics committee. The FID-MRSI was acquired with an acceleration factor 3.5 (20 min, optimal factor determined by assessing CS acceleration) for all three volunteers and were reconstructed with the same regularization as retrospective acceleration case, i.e. *λ* = 3 × 10^−4^. The spectral quality was presented with selected reconstructed spectra from four locations: the cingulate gray matter (GM), the frontal white matter (WM), the caudate nucleus and the temporal GM with the corresponding LCModel fitting.

#### Brain-atlas regional analysis

To illustrate the metabolite contrasts and their relation to the underlying anatomical structures, the 3D metabolite maps were co-registered to an anatomical brain atlas. For each participant, *T*_1_-weighted anatomical scans were segmented into GM and WM compartments by using the computational anatomy toolbox (CAT12; http://www.neuro.uni-jena.de/cat). Masks for the cerebral lobes were generated using the Standard atlas [62] in WFU PickAtlas toolbox (https://www.nitrc.org/projects/wfupickatlas/) and subcortical gray matter structures were automatically segmented using Freesurfer version 6.0.0 [63]. To cope with MRSI partial voluming, a general linear model was employed to fit the metabolite concentration in each anatomical structure (a detailed description of the model is given in the supplementary material). These values were computed for all three volunteers and the mean ratios (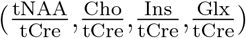) across volunteers were compared with reported values [60, 64–67]. An ANOVA was performed for each metabolite across anatomical structures with the three volunteers data to highlight significant differences in metabolite concentration. Following this test, a multiple comparison was performed to investigate possible significant pairwise differences between individual structures.

## III. RESULTS

The results of the in vivo MRSI data accelerated retrospectively to illustrate the CS performance are shown in fig.2 for the five major metabolites measured in 3T FID-MRSI: N-acetylaspartate + N-acetyl aspartylglutamate (tNAA), creatine + phosphocreatine (tCre), Choline moities metabolites (Cho) (choline, acetylcholine, phosphocholine and glycerophosphocholine), myo-inositol (Ins), glutamate + glutamine (Glx). The cortical layer is visible on tCr and Glx maps and the Cho distribution shows the highest signal intensity in frontal WM and the lowest values in the occipital lobe. These spatial features still remain clear even when data are progressively randomly undersampled in the k-space from 50% to 17% (acceleration factor from 2 to 6). Although all datasets were reconstructed with the same regularization parameter value *λ*, metabolite images gradually lose fine detailed structures and show blurriness as the acceleration factor increases due to the lack of high k-space frequencies remaining sampled in the data. This is highlighted by the generally monotonic rise of RMSE and decrease of SSIM. Thorough simulation of image reconstruction was performed to demonstrate this effect (results are presented in the supplementary material). It is demonstrated that a regularized model with optimal parameter minimizes the reconstruction error, whereas a strong acceleration of the data cannot be reliably reconstructed without important loss of apparent resolution.

**FIG. 2.**
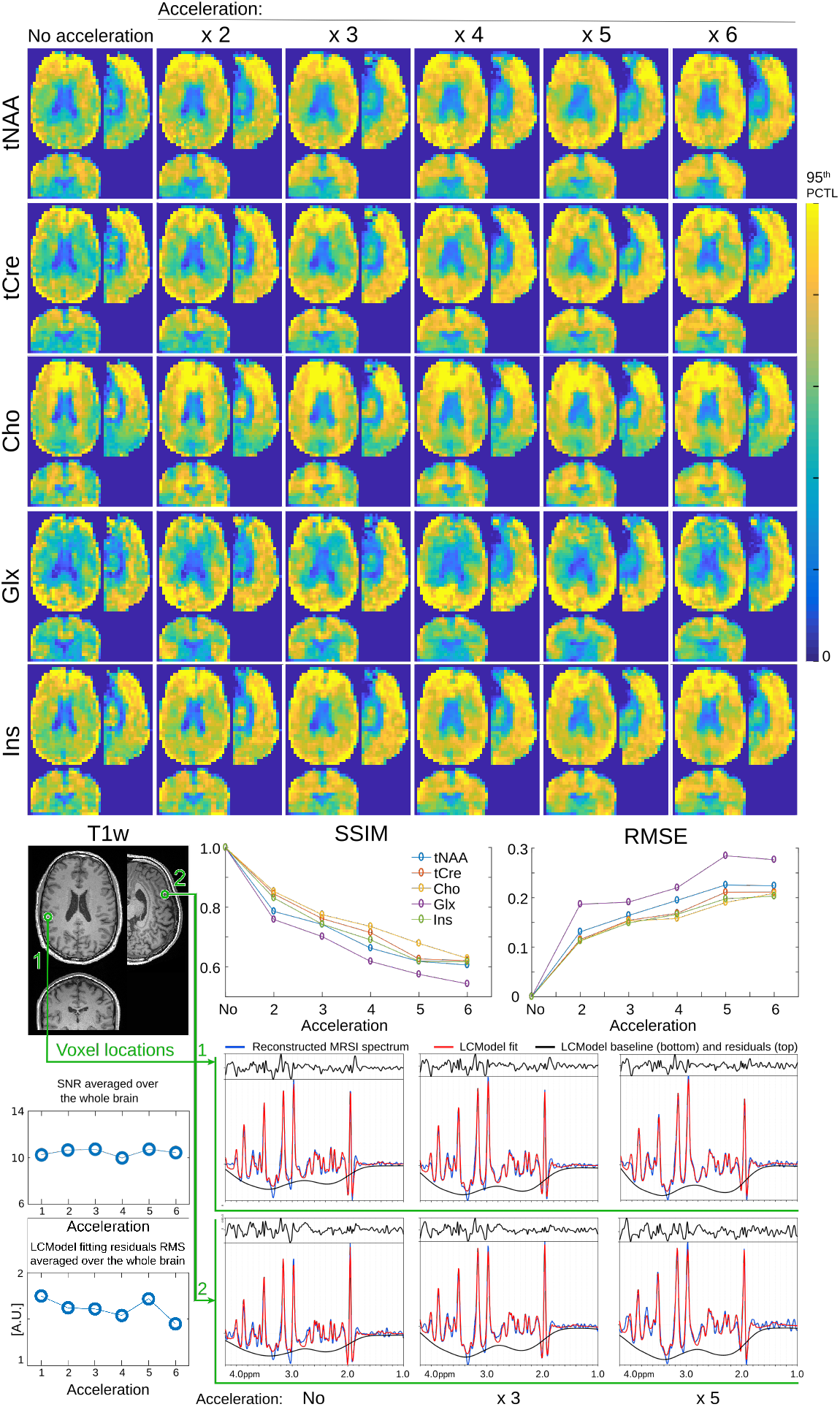
Top, 3D FID-MRSI reconstructed metabolite volumes with retrospective acceleration. The fully sampled acquisition (No acceleration) was acquired in 70 min and acceleration factors correspond to k-space undersampling and reducing acquisition time accordingly (e.g. x3: 24 min, x6: 12 min). The color map was scaled individually for each metabolite range from 0 to the 95th percentile. Bottom, the normalized root mean square error (RMSE) and structural similarity index (SSIM) computed for each metabolite map at all acceleration factors relative to the unaccelerated result. Bottom, sample spectra from two distinct locations are displayed and exhibit very little variation with the acceleration (no, 3, 5). The LCModel fit are shown with the fitting residuals. The root mean square (RMS) of the residuals averaged over the whole brain remains constant with the acceleration (bottom plot).

The RMSE and SSIM of each metabolite volume relative to the ‘no acceleration’ volume show monotonic behaviors close to a linear relation with the acceleration factor in agreement with previous publications [38, 55]. Among all metabolites Glx consistently shows the highest RMSE and lowest SSIM. This is probably due to the lower signal of glutamine and glutamate compared with other metabolites. tNAA also shows higher error and lower similarity in comparison to tCre or Cho. This is possibly relative to the lipid suppression that can affect baseline at 2 ppm and add variability in the tNAA quantification.

Spectra shown at two locations in the bottom of fig.2 exhibit no noticeable change with acceleration. Possible spectral distortion as a result of the acceleration could induce an increase of fitting residuals, however, the residuals root mean square (RMS) average over the brain remains relatively constant.

The full tNAA, tCre, Cho, Ins and Glx maps resulting from the accelerated acquisition (i.e 3.5) are displayed for volunteer 1 in fig.3. The metabolite maps match the distribution features and have similar quality as for the retrospective acceleration by a factor 3 or 4 in fig.2. Volumes of Cho and Glx concentrations which display the sharpest contrasts, and four sample spectra are presented in fig.4 for each of the three volunteers. The concentrations variations observed in the metabolite maps translate into visual spectral variations. The frontal WM spectrum shows a clear higher Cho peak with respect to the other three locations, and the cingulate GM and temporal GM spectra present markedly higher Glx signal than the frontal WM and caudate. These visual landmarks are noticeable for all three volunteers. The metabolite concentration ratio maps of the same measurements are shown in supplementary Fig.S9. In addition, CRLB and SNR maps of these results are presented in supplementary Fig.S10 but should be interpreted with caution. Indeed, their accuracy might be affected by the reconstruction process as described in the Methods. To validate the quantification in I.U. with respect to the water reference signal, MRSI data were measured, reconstructed and quantified on a homogeneous metabolite solution phantom (MRS Braino phantom, GE Medical Systems, Milwaukee, WI, USA). The resulting metabolite map (supplementary Fig.S16) exhibit homogeneity over the whole phantom.

**FIG. 3.**
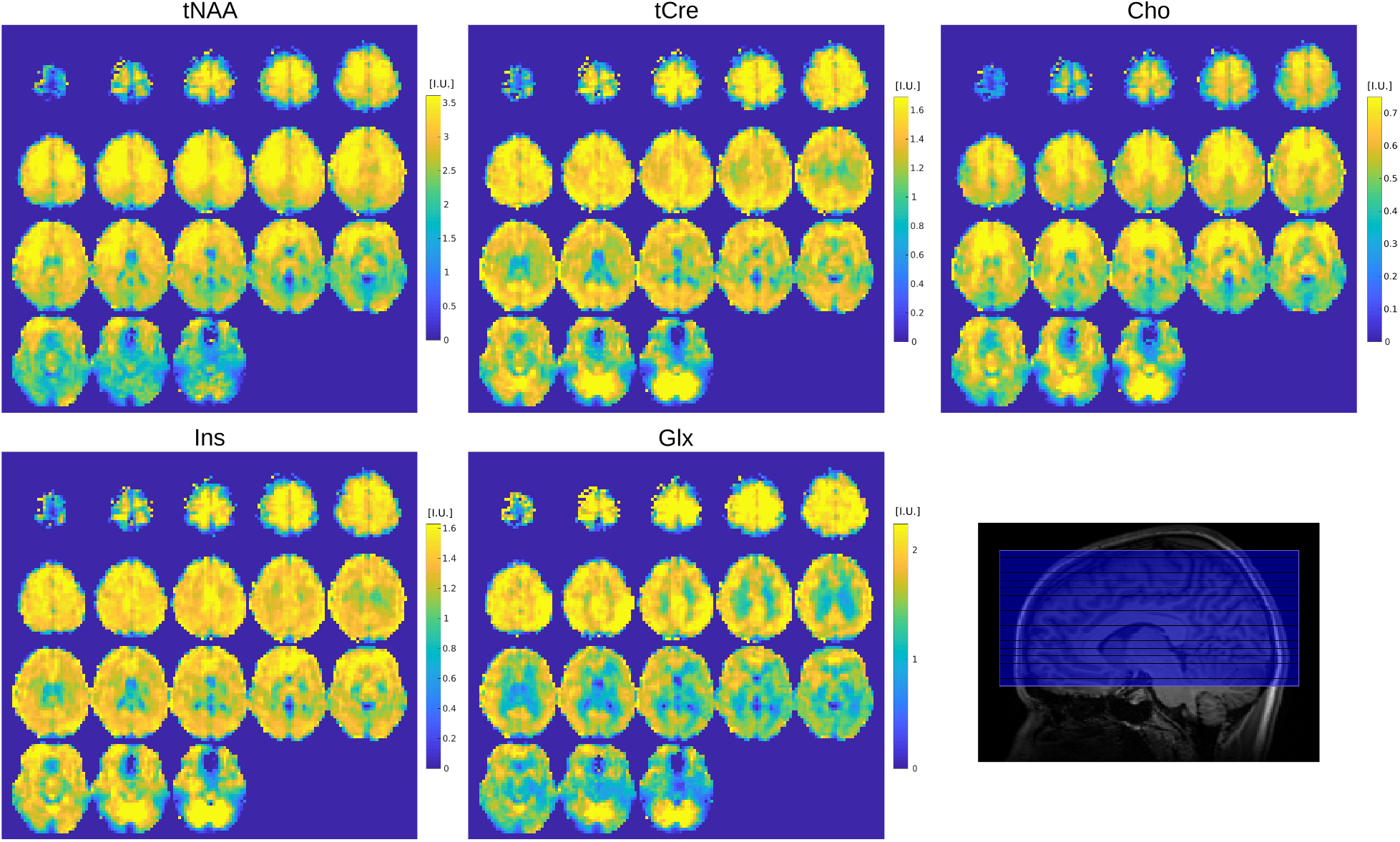
CS-SENSE-LR 3D FID-MRSI measured on a healthy volunteer (volunteer 1) with 5 mm isotropic resolution in 20 min with acceleration factor 3.5, resulted in tNAA, tCre, Cho, Ins and Glx maps. Color scale for each map is given in institutional units. Sagittal *T*_1_-weighted image (bottom right) show the location of the excitation slab (blue overlay).

**FIG. 4.**
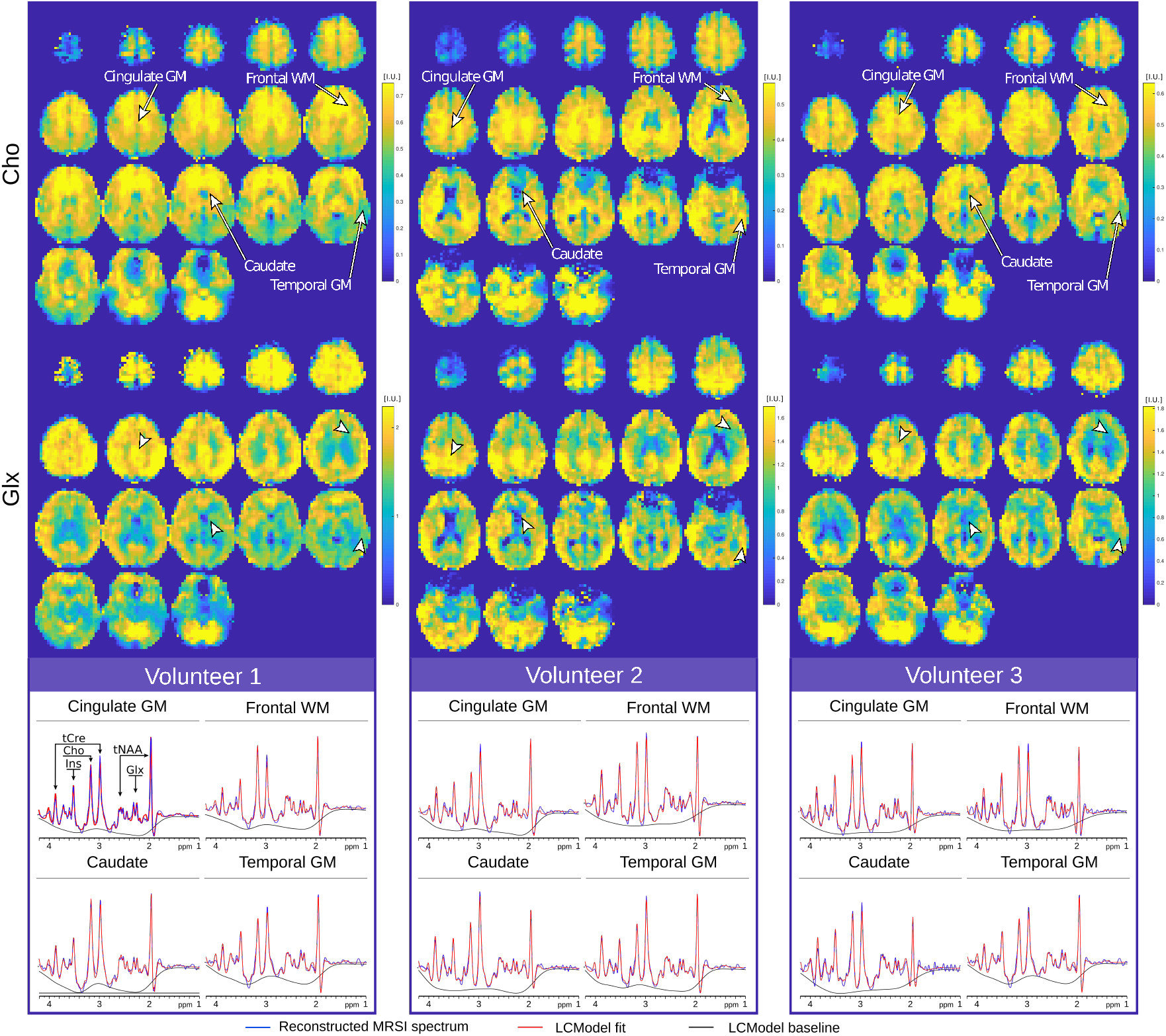
Comparison of high-contrast Cho and Glx 3D maps measured with CS-SENSE-LR FID-MRSI on three healthy volunteers (volunteer 1, 2 and 3) with 5 mm isotropic resolution in 20 min (acceleration factor 3.5). Color scale is given in institutional units for the respective metabolite concentration. Samples spectra originating from four distinct locations are shown for each volunteer. Cho and Glx signal amplitude in the spectra match the metabolite distribution observable on the maps. Metabolite ratio maps of the same volunteers and the corresponding Cramer-Rao lower bound (CRLB) and SNR maps estimated by LCModel are shown in supplementary Fig.S10.

The spectra shown in fig.4 and fig.2 exhibit high SNR as a result of the low-rank constraint and a narrow linewidth due to the small voxel size as well as the *B*_0_ fieldmap correction included in the CS-SENSE-LR model. Mean whole-brain SNR estimated by LCModel was 14.2, 12.7 and 12.6 and the mean spectral full width at half maximum was 5.67 Hz, 5.46 Hz and 5.19 Hz for volunteers 1 to 3, respectively. The baseline distortion due to the lipid suppression by orthogonality is visible as a pit at 2 ppm forming a ‘W’ shape with the NAA peak. Nevertheless, the distortion is well fitted by the inverse broad peak added to the basis and the resulting baseline is smooth. The metabolite maps remain sensitive to *B*_0_-field inhomogeneity that results in local linewidth increase. This effect is visible in the lower sections of the metabolite maps, *B*_0_ fieldmaps and linewidth maps (fig.4, supplementary Fig.S12, supplementary Fig.S11).

The anatomical segmentation of 3D metabolite maps for volunteer 1 is illustrated in fig.5. Concentrations of each anatomical region are reported in fig.6 and illustrate the consistency of the anatomical contrasts for all volunteers with the distribution of metabolite concentrations following similar patterns. For tNAA, cortical concentrations tend to be lower in temporal lobes while subcortical concentrations are relatively high in the thalamus. Similarly, tCre concentrations are lower in temporal lobes than in other cortical regions. There is also a systematic difference in tCre between GM and WM. For Cho, plots show a lower concentration in GM than in WM as well as a high concentration in the brain stem, hippocampus and amygdala. In contrast to Cho, Glx exhibits a consistent and markedly higher concentration in GM than in WM. The distribution of Ins concentrations follows a GM-WM contrast similar to tCre. Concentrations in the cerebellum are marked by high concentration of tCre, Glx and Ins. The ratios of metabolite concentrations in each anatomical region, averaged across subjects are presented in fig.7 Values from left and right hemisphere were combined to facilitate comparison with results from the literature [60, 64–67]. Globally, results of tNAA/tCre, Cho/tCre and Glx/tCre match the previously reported ratio values with differences that are in the range of what has been previously published, despite tNAA/tCre, Cho/tCre values are in the highest reported values and Glx/tCre in the lowest. The GM-WM contrast that is reported in the literature for all metabolite ratios, is also observed in the present study. Ins/tCre values are systematically larger than the literature results although the GM-WM contrast here is comparable. Results of the statistical analyses of differences across brain regions are presented in supplementary Fig.S15. The ANOVA analysis exhibited a significant difference across regions in all the metabolites (p-value given in the title of each figure). The results of the multiple comparison testing are presented as matrices for each metabolite. Significant differences (corrected *p <* 0.05) are consistent with the qualitative results described above. Cerebellum concentration is significantly higher compared to various other structures for tCre, Cho and Glx. The cortical GM regions are significantly different from WM and deep GM structures regarding Glx and tCre. Ins and tNAA show fewer significant differences between deep GM structures or between deep GM structures and lobe values.

**FIG. 5.**
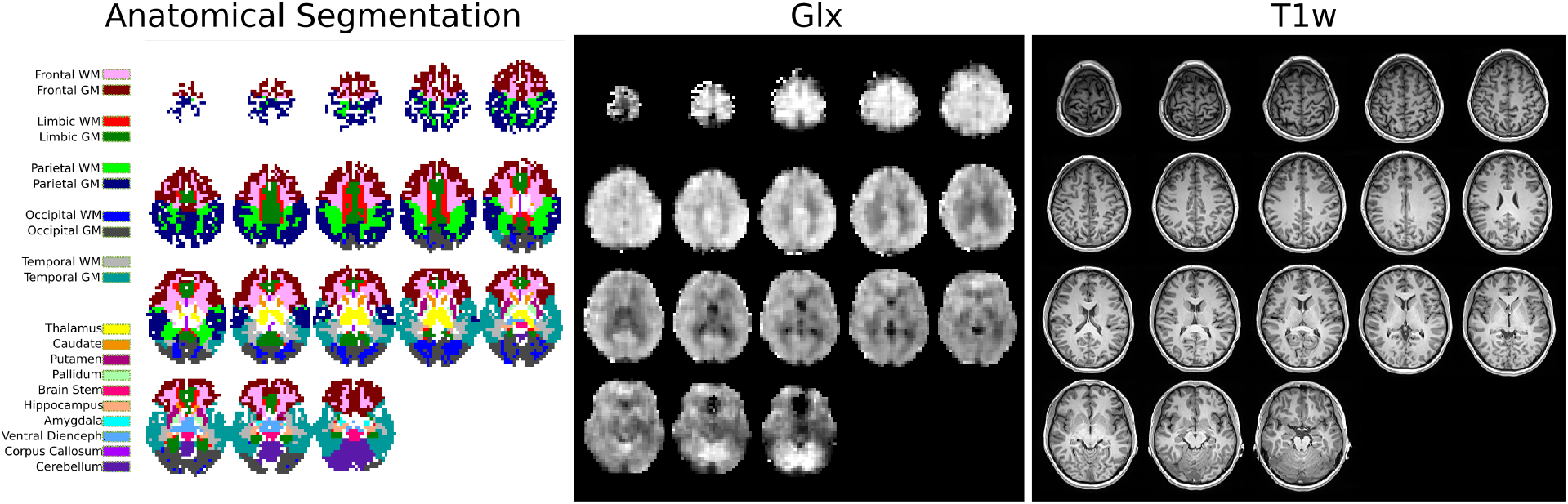
Left, the anatomical atlas registered to volunteer 1 is shown with voxel labeling corresponding to the dominant partial-volume. Center, Glx concentration 3D map is shown for qualitative comparison in a gray scale and, right, the MPRAGE *T*_1_-weighted images used to segment the volunteer anatomic parts.

**FIG. 6.**
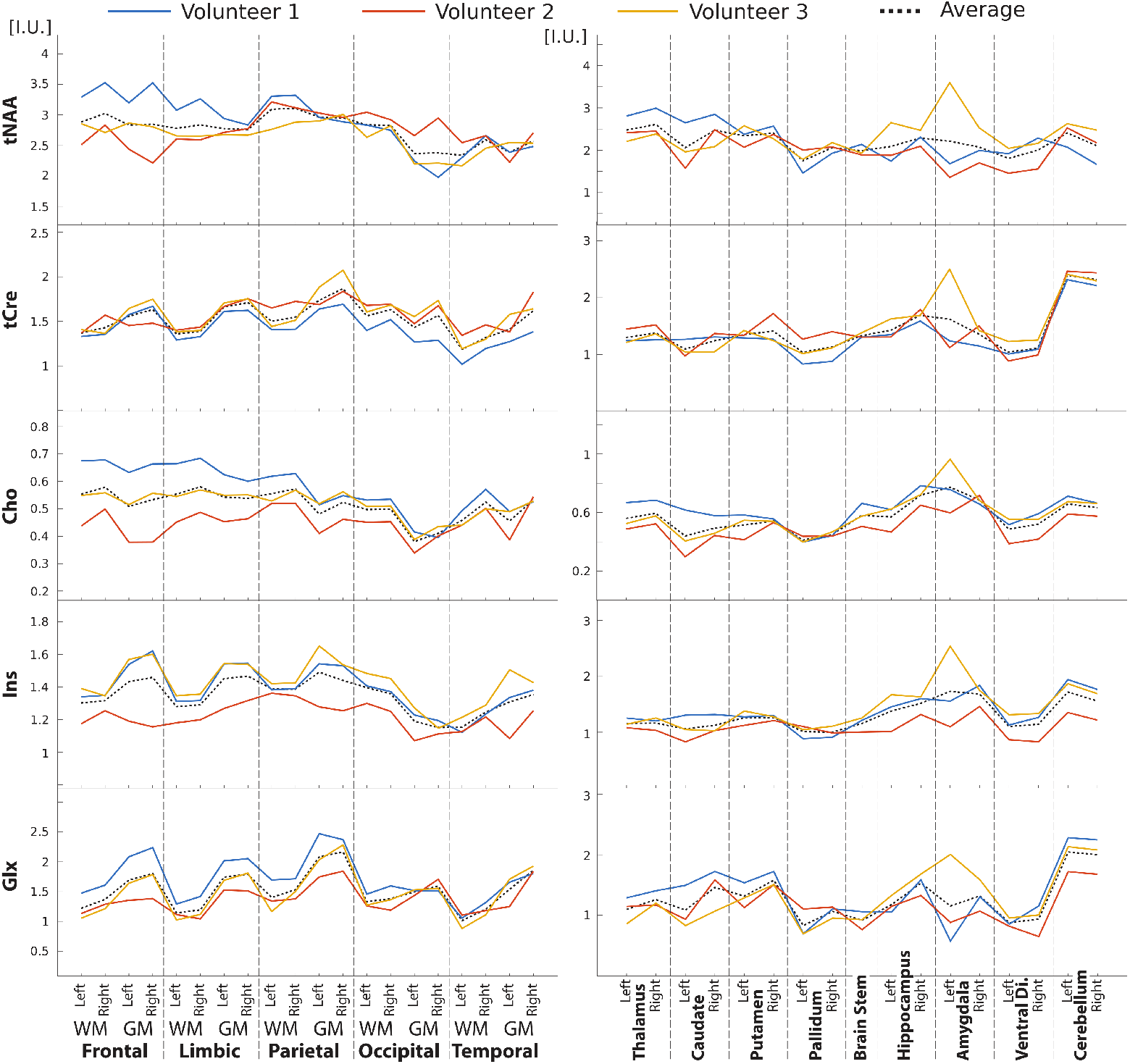
Atlas metabolite content across three healthy volunteers based on anatomical segmentation (fig.5). Lines correspond to the metabolite concentration estimated for all anatomical area for each of the three volunteers. The dashed line represents the mean values across the three volunteers. Metabolite levels are expressed in institutional units. Left plots represent relative concentrations in WM or GM in each cerebral lobe while right plots show relative concentrations in the deep GM.

**FIG. 7.**
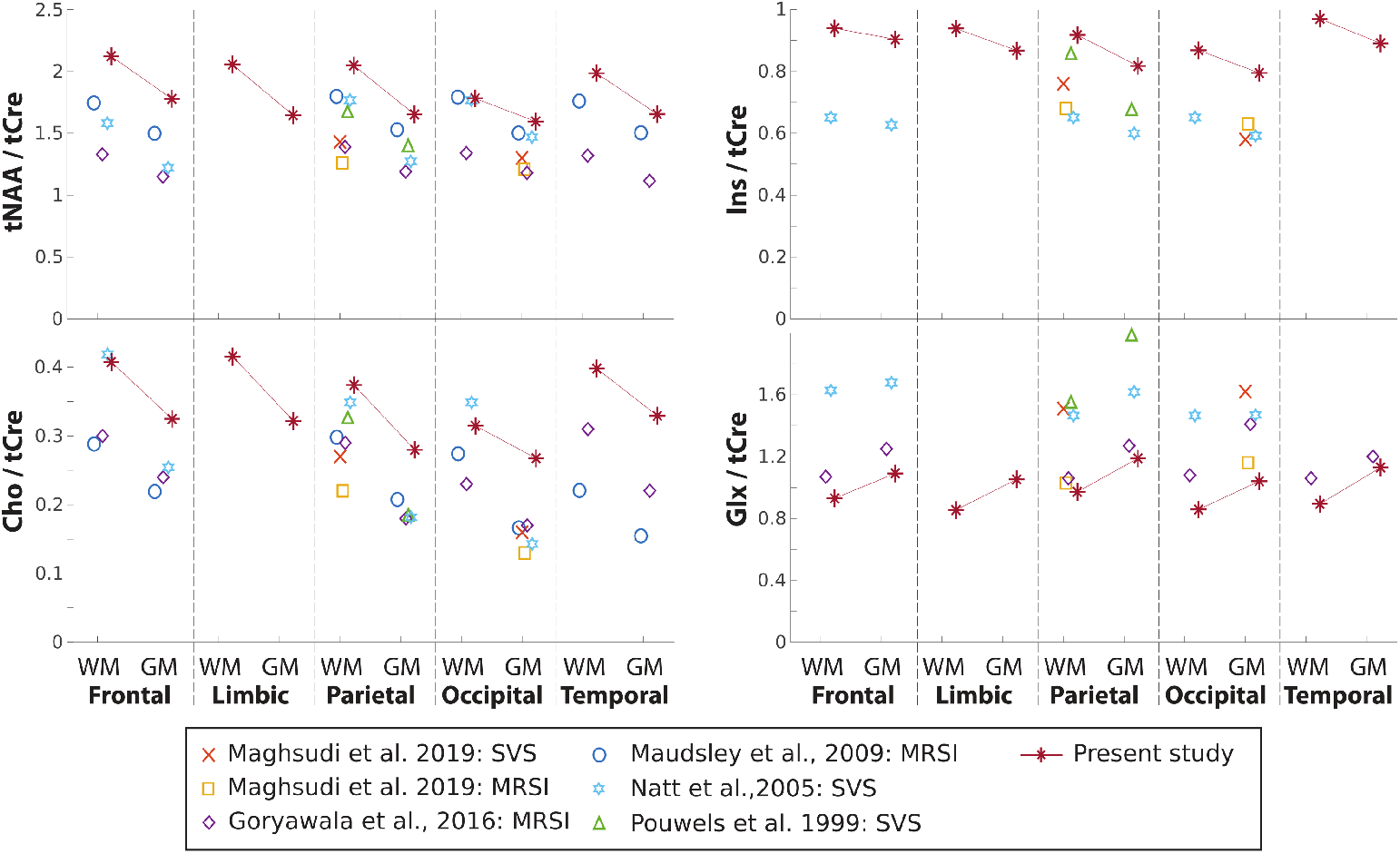
Comparison of the atlas metabolite-ratio content to previously published results. Line-connected points correspond to the metabolite average ratio from volunteers 1 to 3 (individual data presented in fig.6). The legend indicates the corresponding study and the type of acquisition: whole-brain MRSI or sing voxel spectroscopy (SVS)

## IV. DISCUSSION

There is a strong need for the whole-brain mapping of the metabolite distributions in fine details and within an acceptable time frame to allow its inclusion in scanning protocols running on regular 3T MR systems. Here, for the first time, we used the combination of FID-MRSI with CS-SENSE-LR to reconstruct high-contrast and high-resolution 3D metabolite volumes following an acquisition limited to 20 min on a clinical 3T MRI.

The whole-brain approach coupled with the high-resolution (i.e. 5 mm isotropic) of the reconstructed volumes allowed a quantitative analysis of the brain metabolites in all brain regions. Results from the three volunteers showed the same patterns of contrast between cerebral lobes, subcortical structures and GM/WM segments for each metabolite, illustrating consistency between measures.

### Acceleration performance, artifacts and limitations

The performance of the CS-SENSE-LR model in reconstructing undersampled dataset as depicted in fig.2 is demonstrated by the retrospective acceleration (or k-space undersampling) of a fully sampled 3D FID-MRSI. The random k-space spheroid sampling following a radius^−1^ distribution rule only affects the resulting spatial metabolite distributions without downgrading the spectral quality. Indeed, the variable density undersampling scheme mostly removes acquisition points at the periphery of the k-space while preserving the center that contains a majority of the signal. Consequently, spectral SNR is practically preserved using this acceleration scheme. This reconstruction property was highlighted by the qualitative and quantitative analysis of spectra after the retrospective acceleration in fig.2 and was also reported in a previous publication [38]. When the acceleration factor is increased, fine-scale details gradually disappear. As stated above, this artifact is caused by the undersampling of the high k-space frequencies [43, 44]. The same effect could be reproduced with simulation data on two reference images, supplied as supplementary material (fig.S3, S4, S7 and S8). For high acceleration factors images show erosion of small structures even when the regularization parameter value is optimal. This non-uniform effect might differ for some fine-scale features or large image elements [44]. As illustrated by the increase of RSME and decrease of SSIM, this effect on the metabolite maps is already present at small acceleration factor and becomes increasingly visible. The aim of the retrospective analysis shown in fig.2 is to find the right balance between an acceptable loss in metabolite map quality and a substantial gain in acquisition time. Based on the present results, an acceleration factor of 3.5 was found to be optimal. A higher acceleration factor is accompanied by a loss of apparent resolution and illustrates that the method allows only low to moderate acceleration in comparison to other acceleration methods [13, 14, 24]. Further details on the determination of the regularization parameter *λ* and the rank *K* for the reconstruction, and discussion of the spectral quality can be found in the supplementary material.

The acceleration factor 3.5 of our protocol provides a drastically shorter acquisition (20 min) in comparison to the fully sampled protocol (70 min). However, the MRSI acquisition, even accelerated, remains markedly long and is therefore susceptible to subject motion. While no motion correction or compensation were used in this study, the implementation of an interleaved imaging-based volumetric navigators could be expected to further improve the measurement accuracy and prevent possible motion artefact as shown in [40].

Although the lipid suppression by orthogonality performed as pre-processing efficiently removes any lipid signal from the brain spectra, it creates a specific distortion of the baseline at 2 ppm particularly visible in the spectra of fig.2 and less marked in fig.4. The distortion may strongly affect LCModel fitting performance that compensate the pit by an unrealistic 1st order phase correction impacting the quantification stability. We partially solved the issue by introducing a broad inverse peak in the LCModel basis that allows the distortion pit to be fitted as a negative peak while preserving the spectral phase and baseline estimation. While not being the scope of this article, this coping technique might alter the quantification of NAA and NAAG and further investigations are necessary to ascertain the fitting of these metabolites. The spectral distortion is also more present in part of the FOV with *B*_0_-field inhomogeneity. In these regions with increased linewidth, the orthogonality between metabolite and lipid signal is less valid and a stronger baseline pit is created around 2 ppm. See for instance the lower part of the brain in fig.3 where tNAA and Glx signal are decreased with the corresponding *B*_0_-fieldmap and linewidth in supplementary fig.S11 and S12. Also any metabolite signal located between 1.8 and 1.2 ppm like lactate and alanine peaks, is probably partially removed in the process and their quantification remains inaccurate as suggested also in [68, 69]. The lipid suppression remains nevertheless a necessary step of the pipeline that must be performed before the model reconstruction. Indeed, the low-rank reconstruction provides a solution containing the dominant components in the signal. If the lipid suppression is omitted or not performed before the reconstruction step, most of the model components are contaminated by lipid signal, which overcomes the metabolite signal by several order of magnitude, and a correct metabolite mapping is then impossible. Illustration of the lipid contamination for reduced lipid suppression can be found in supplementary Fig.S20.

Furthermore, in order to prevent this baseline distortion, the lipid suppression by orthogonality could be replaced by other sequence-based lipid removal methods [70] such as saturation radio-frequency pulses [42], inversion recovery [71] or pulsed magnetic field gradients [72] that may better preserve the metabolite signal around 2 ppm. The CS-SENSE-LR acquisition and reconstruction pipeline could be directly applied to these techniques. An additional limitation of the lipid suppression by orthogonality performed during pre-processing is the removal of the macromolecule (MM) signal. In contrast to the metabolite signal, the MM signal does not appear to be spectral orthogonal to skull lipid signal contrarily to the metabolite signal. The MM signal is identified as lipid and is removed during the lipid suppression step. Measuring MM with the presented technique would require the replacement of the lipid suppression by orthogonality with a sequence based lipid suppression method [42, 72].

The use of the FID-MRSI sequence enables the acquisition of MRSI data with an ultra-short TE. This could be advantageous because a TE*<*1ms prevents essentially any *T*_2_ relaxation weighting in the measured data. In contrast, the short TR employed with FID-MRSI in the present study may result in *T*_1_-weighting of the metabolite signal. Indeed, the steady-state magnetization of each metabolite consecutive to the reduced TR depends on the respective metabolite *T*_1_ value. Following previously published results [73, 74], brain metabolite *T*_1_ values are expected to range from 1000 to 1400 ms, and the steady-state magnetization may therefore vary by ±7% (see supplementary Fig.S18). Nevertheless, this possible *T*_1_-weighting represents a signal variation smaller than the contrast observed in the metabolite maps (see e.g. fig.6).

## Results and the literature

The 3D metabolite distributions observed over the whole brain in fig.2, 3 and 4 contains features and contrast described in III that are in agreement with previous publications. Thus, the distribution of tNAA, tCr, Cho exhibit similar patterns as published in [65, 75] and Glx in [10, 40, 66]. Ratio results in the present study are comparable with those reported in the literature. In particular, the GM-WM contrast previously observed across healthy brains in most metabolites is reproduced by the present study. Differences are observed in the Ins/tCre values that are systematically higher in our results. Also Glx/tCre seems systematically low, although considering the variability across previously reported results, it remains in good agreement, particularly with the data from Goryawala *et al*. [66]. These discrepancies could be caused by the difference in acquisition sequence. All previously reported results were acquired with a spin echo sequence and with a significantly longer TE (*>*20 ms). As described above, the difference in TE might greatly influence the quantification of some J-coupled metabolites. To date, there is no other FID-MRSI results reported on atlas permitting a driect comparison.

The magnitude and the consistency of these contrasts are emphasized by the statistical analysis of the data across the three volunteers. Several significant differences were found between regions exhibiting high contrast in the metabolite maps (see supplementary Fig.S15).

The consistency of the segmentation results across volunteers shown in fig.6 illustrates qualitatively the sensitivity of the technique but an actual reproducibility study would be useful to assess the variability and to disentangle intersubject physiological difference from inter scan methodological variability. Based on our presented results, an application to SSE with undersampled trajectory should be considered and would yield even shorter acquisition time for equivalent or higher resolution. The acquisition-reconstruction method proposed in this study can be compared to other published work on full brain MRSI at 3T [24, 39, 40, 76]. The CS-SENSE-LR model reconstruction requires additional hours compared to other methods and thus may be an obvious limitation in its application. However, hardware improvement, software optimization, parallelization and algorithm development could dramatically reduce the processing time.

## Conclusion

To conclude, a novel acquisition-reconstruction scheme, combining FID-MRSI with CS-SENSE-LR, enables 3D spectroscopic imaging of the whole human brain in high-resolution on a 3T system. The reconstructed metabolite volumes showed high anatomical contrast and high levels of features in 5 mm isotropic resolution. The resulting spectral quality demonstrated the efficiency of the model reconstruction for SNR enhancement and *B*_0_ field map correction. Acceleration by random k-space undersampling allowed a dramatic reduction of the acquisition time from 70 to 20 min which makes its implementation in clinical or research protocols feasible. As proof of concept, metabolite volumes from three volunteers were segmented into anatomical lobes and substructures. The resulting contrast observed quantitatively is the counterpart of the metabolite features visible on 3D metabolite maps and are consistent through all three volunteers.

## V. ACKNOWLEDGMENTS

PK is supported by a fellowship from the Adrian and Simone Frutiger Foundation.

