## supplementary material for "Whole-Brain High-Resolution Metabolite Mapping with 3D Compressed-Sensing-SENSE-LowRank ^1^H FID-MRSI"

Further details about the methods and results of the main manuscript are presented in this document.

### I. TGV RECONSTRUCTION DEMONSTRATION

In this section, we aim to demonstrate the accuracy and validity of image reconstruction with total generalized variation (TGV) constraint. Two synthetic $128 \times 128$ images were used for benchmarking of the TGV reconstruction that are presented in fig.S2. The first image is a binary pattern aiming to illustrate effect of reconstruction on edges with small structures with strong contrast. The second image is more realistic and represents a T1-weighted MRI image of the human brain. The raw data corresponding to these two images were created by Fourier transform and the Fourier domain (k-space) data were masked to match a specific elliptical sampling pattern with or without undersampling (see k-space masks in fig.S1). Following notation of the main manuscript, the image $\rho(\mathbf{r})$ solution to the inverse problem is reconstructed by minimization of the TGV-constrained problem

$$\arg\min_{\rho(\mathbf{r})} \; \|\mathbf{s} - \mathcal{F}(\rho(\mathbf{r}))\|_2^2 + \lambda \mathrm{TGV}^2\{\rho(\mathbf{r})\} \tag{1}$$

with $\mathbf{s}$ the synthetic image k-space data described above, $\mathcal{F}$ the encoding operator (including Fourier transform and k-space masking). The reconstruction was performed for the regularization parameter $\lambda$ ranging exponentially from $10^{-6}$ to 1 and for random undersampling of 100%, 50%, 33%, 25%, 20% and 17% corresponding respectively to acceleration factors (A.F.) of 1 (no undersampling), 2, 3, 4, 5 and 6.

The proposed problem consisting of the brain and pattern image reconstruction with TGV regularization and undersampled k-space, is equivalent to the problem of retrieving the MRSI spatial components $U_n(\mathbf{r})$ presented in the main manuscript. Indeed the MRSI data undersampling were only performed spatially and a partial spatial-spectral separation was assumed leading to a reconstruction problem of spatial components $U_n(\mathbf{r})$ equivalent to an image reconstruction problem. Therefore, the conclusion about the effect of the k-space undersampling and TGV regularization of this proposed problem can be translated into effects over the resulting metabolite concentration maps and their respective contrast or artifact. The brain and pattern image reconstruction problem is employed here for the demonstration because the artifact and undersampling effect are easier to identify and interpret on the presented images than on reconstructed MRSI dataset.

#### A. TGV regularization minimizes the reconstruction error

In this part, it is demonstrated that the TGV regularization minimizes the reconstruction error with respect to the reference images. This error is quantified as the normalized root-mean-square error ($NRMSE = \frac{\sum_{\mathbf{r}} |\rho_{recon.}(\mathbf{r}) - \rho_{ref.}(\mathbf{r})|}{\sum_{\mathbf{r}} |\rho_{ref.}(\mathbf{r})|}$). Results of the reconstruction are presented for a subset of $\lambda$ and few A.F. for the brain image (fig.S3) and the pattern image (fig.S4). In both figures, the most left column represents the reconstruction results with a very low TGV regularization, equivalent to a reconstruction without TGV. The solution to this case is given by the Penrose pseudo-inverse operator and simplifies into the usual inverse Fourier transform of the k-space data. These results show minimum error in absence of undersampling (but non-zero due to elliptical sampling) and exhibit strong alteration with increasing acceleration as consequence of the k-space undersampling and resulting in noise-like aliasing. As $\lambda$ increases, NRMSE decreases first and image quality improves in all cases without exhibiting particular smoothness (although loss of contrast is visible for strongly undersampled data. This is discussed lower in details with analysis of the image profiles). As $\lambda$ is further increased, NMRSE increases and results show loss of edges and over-smoothening, the sign of over-regularization. This is particulalrly obvious in the most right column in fig.S3 and fig.S4. The optimal $\lambda$ parameter is determined by the minimum NRMSE in fig.S5: $\lambda_{opt} \approx 1.5 \cdot 10^{-3}$ for the pattern image and $\lambda_{opt} \approx 5 \cdot 10^{-4}$ for the brain image. NRMSE can however only be computed when reference data are accessible. This is not the case for in-vivo data and often the L-curve criteria is employed as surrogate technique to determine $\lambda_{opt}$[1]. L-curves are shown here for the pattern and brain reconstructions in fig.S6. Usual criterion to determine $\lambda_{opt}$ is the point of maximal curvature. It is however notcieable that this technique tends to underestimate the optimal TGV regularization parameter by a factor 3 (maximal curvature being around $\lambda = 4 \cdot 10^{-4}$ for the pattern image and $2 \cdot 10^{-4}$ for the brain image).

These results illustrate that an optimal regularization permits to improve the reconstruction of randomly undersampled data without necessary introducing smoothness in the image. With the optimal $\lambda_{opt}$, the regularization guides the reconstruction of the underdetermined inverse problem towards a solution spasifying edges and second order derivatives, minimizing the presence of k-space undersampling artifact. More technical aspects and demonstrations can be found in the compressed-sensing seminal papers [2, 3]. Similar results were obtained from a quantitative analysis on simulated MRSI data in [4].

#### B. Contrast and edge preservation

The effect of the TGV regularization and the undersampling on the contrast and the image edges is analyzed in further details here. In fig.S7 and fig.S8, a profile chosen in both the brain and pattern images is displayed. In fig.S7 left plot, the data presented for 50% undersampling (A.F.=2) demonstrate that the profile can be fully retrieved with the regularization parameter chosen in the vicinity of the optimal value ($3 \cdot 10^{-4} \leq \lambda \leq 3 \cdot 10^{-3}$). In the under-regularized regime ($\lambda < 3 \cdot 10^{-4}$), pseudo-random aliasing due to undersampling is present over the whole profile (particularly visible in the first points of the profile with ground-truth value being 0). With over-regularized solution ($\lambda > 3 \cdot 10^{-2}$), the profile is clearly smoothened with strong attenuation of the edges. The right plot in fig.S7 shows the resulting profile for all k-space underampling factors but reconstructed with $\lambda_{opt}$. For undersampling $\geq 0.33$ (A.F.$\leq 3$), the profile is practically fully retrieved by the reconstruction with the exception of isolated voxel that is attenuated for A.F.$= 3$. For greater A.F. the profile suffers significant loss of contrast and edges are strongly softened. Similar observations can be made in fig.S8 with the brain image although there are less marked edges in the original reference image. Here we demonstrated that up to a certain A.F. with k-space undersampling $\geq 0.33$, the original image can be faithfully retrieved by the reconstruction regularized with the optimal parameter $\lambda_{opt}$ without increase of NRMSE, loss of edges or contrast. For sampling $< 0.33$, even with the optimal parameter $\lambda_{opt}$, the reconstructed image suffers from significant loss of image features and displays an apparent smoothness due to the loss of small details. This effect can be understood by observing the undersampling masks for strong acceleration in fig.S1. For high undersampling, there is practically no point sampled at the periphery of the k-space. These points however contain the high spatial-frequency informations of the original image and therefore are necessary to reconstruct small details and contrasts of the image. If too much undersampling is performed in the high-frequency k-space points, the reconstruction is unable to retrieve the precise image even with optimal regularization. The apparent smoothness or loss of details cannot be translated into a global increase in the effective voxel size due to the non-linearity of the reconstruction process. The resulting smoothness would differ from one part of the image to another based on the content. It cannot be descibed as a global smoothing process (such as moving average or a filter).

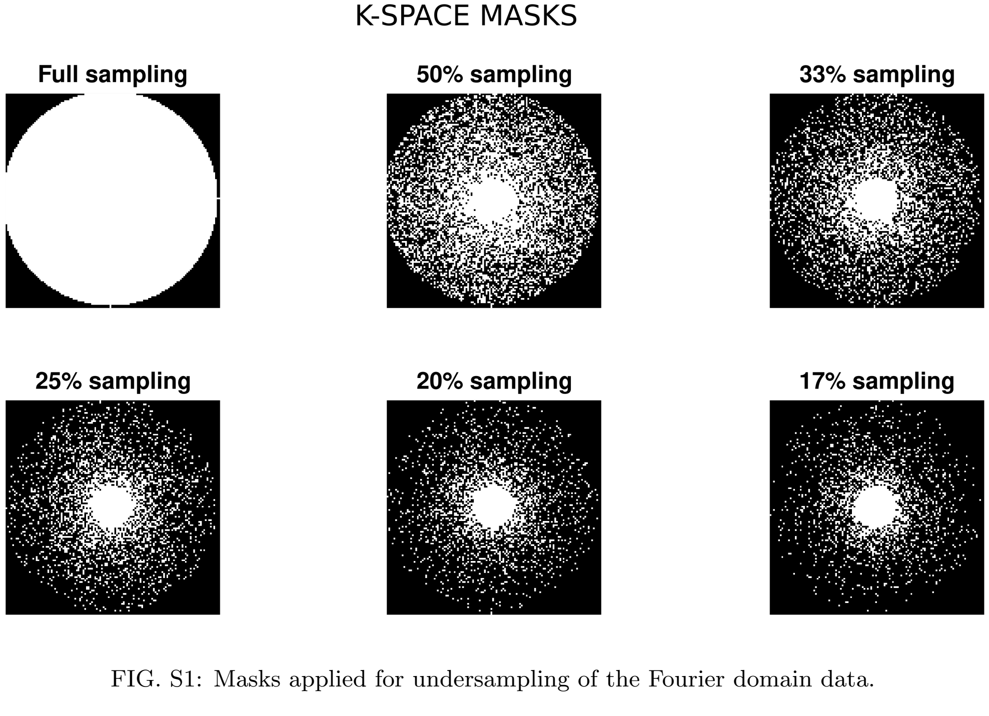


FIG. S1: Masks applied for undersampling of the Fourier domain data.

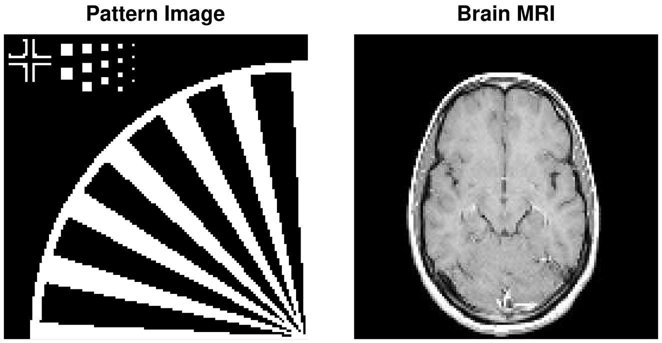


FIG. S2: Images used to assess the reconstruction performance: left, a simple binary pattern to highlight effects on edges and voxel size. Right, a T1-weighted human brain MRI image.

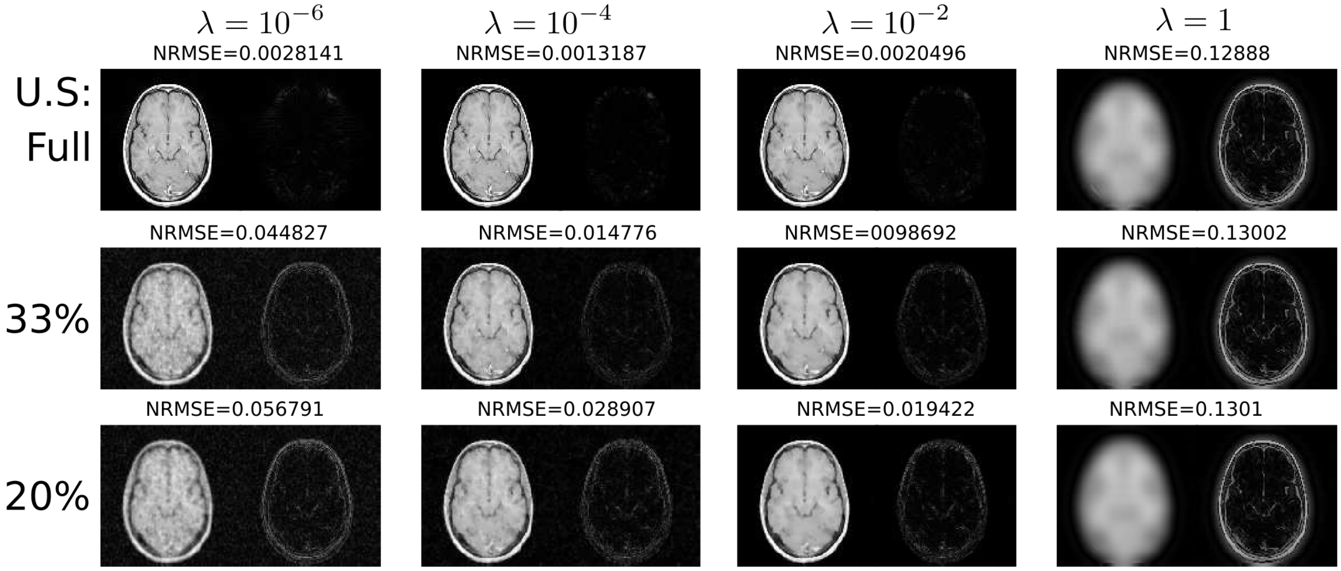


FIG. S3: Reconstructed brain images (left) and the error to the original image (fig.S2) (right) for varying TGV regularization ($\lambda$) and k-space undersampling (U.S).

### II. DETAILS ON THE BRAIN-ATLAS REGIONAL ANALYSIS

To illustrate the metabolite contrasts and their relation to the underlying anatomical structures, the 3D metabolite maps were co-registered to an anatomical brain atlas. For each participant, $T_1$-weighted anatomical scans were segmented into gray and white matter compartments by using the computational anatomy toolbox (CAT12; http://www.neuro.uni- jena.de/cat) as implemented in statistical parametric mapping (SPM12) software (https://www.fil.ion.ucl.ac.uk/spm/software/spm12/) running in Matlab R2018b (The MathWorks, Inc., Natick, Massachusetts, US). Native-space gray matter images were spatially normalized to the DARTEL template in MNI standard space created from 555 healthy control subjects from the IXI-database (http://www.brain-development.org). Masks for the cerebral lobes were generated using the Standard atlas [5] in WFU PickAtlas toolbox (https://www.nitrc.org/projects/wfu_pickatlas/) and spatially transformed in participants' native space by applying the inverse of the transformation matrix generated during the spatial normalization step. Subcortical gray matter structures were automatically segmented using Freesurfer version 6.0.0 [6]. The resulting anatomical atlases were first registered to the MRSI orientation (main manuscript fig.5). The MRSI voxel size leads to partial voluming and a single MRSI voxel might contain contributions from several anatomical structure simultaneously. To cope with this partial volume effect and accurately estimate the concentration in each anatomical structure, a general linear

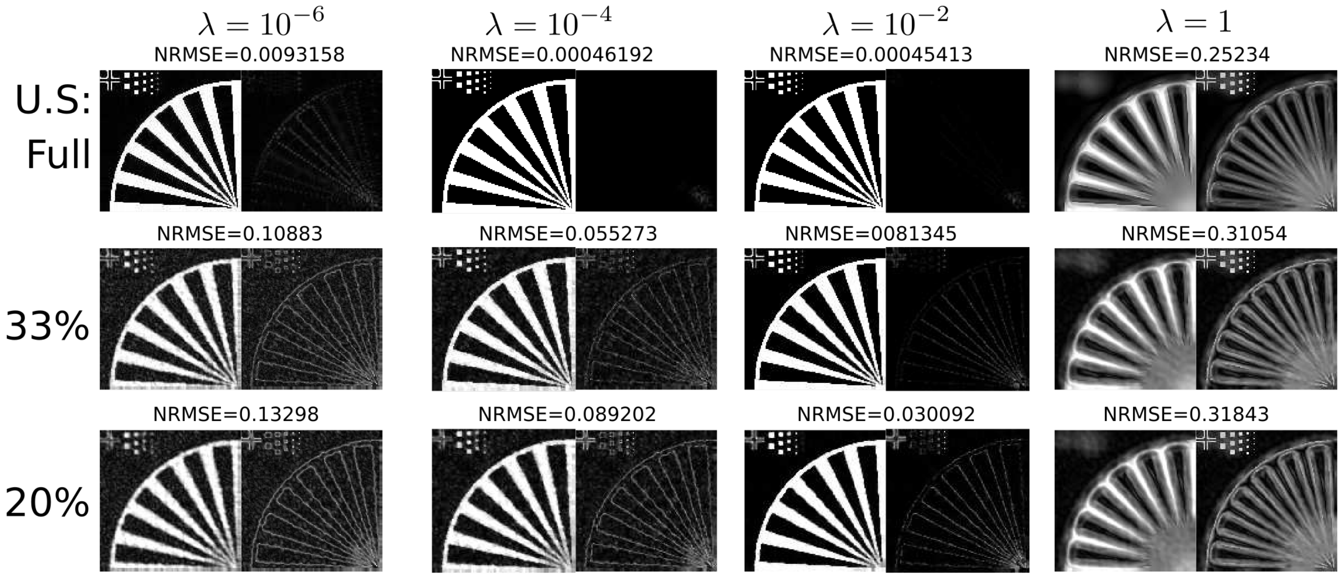


FIG. S4: Reconstructed pattern images (left) and the error to the original image (fig.S2)(right) for varying TGV regularization ($\lambda$) and k-space undersampling (U.S).

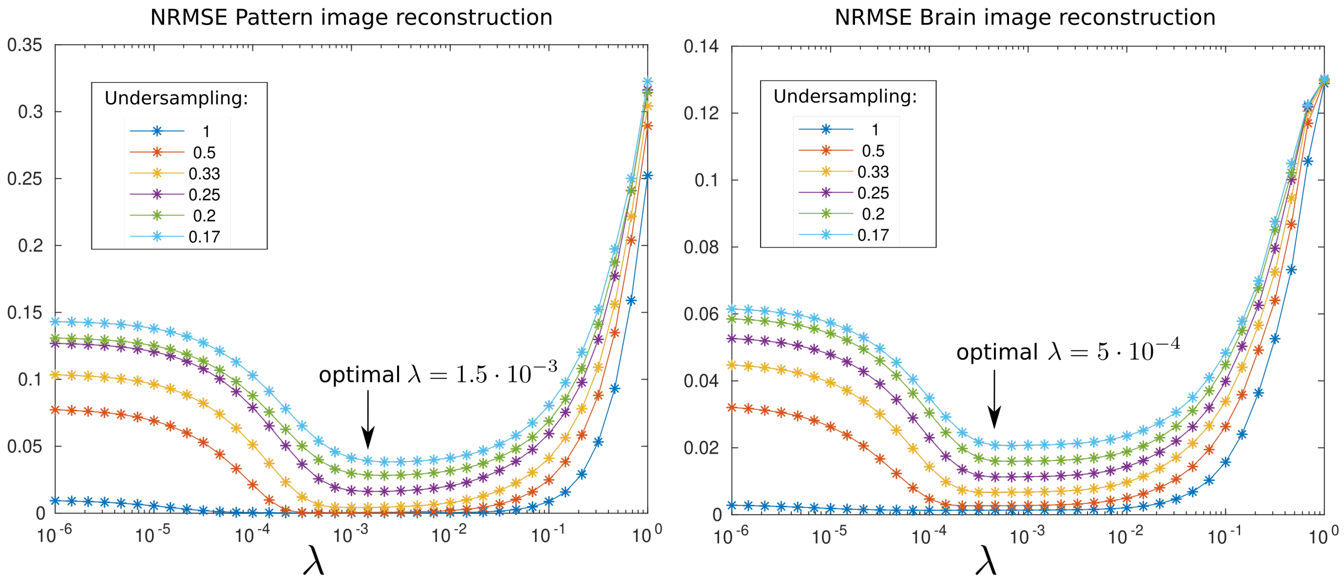


FIG. S5: Normalized root-mean-square error of the reconstructed pattern and brain images as function of the TGV regularitzation parameter ($\lambda$) and for several undersampling of the k-space.

model was employed for each metabolite [7]:

$$\mathbf{Y} = \sum_i \beta_i \mathbf{X_i} + \beta_0 + \epsilon \tag{2}$$

where $\mathbf{Y}$ is a vector made of the metabolite concentration in all voxels, $\mathbf{X_i}$ is a vector containing all partial-volume estimate (a value between 0 and 1) with $i$ the index of the anatomical structures presented in main manuscript fig.5, $\beta_0$ is the concentration intercept and $\epsilon$ represents normally distributed residual. Following the fit of the general linear model, the metabolite concentration in each anatomical structure corrected for the partial-volume is given by $\beta_i + \beta_0$.

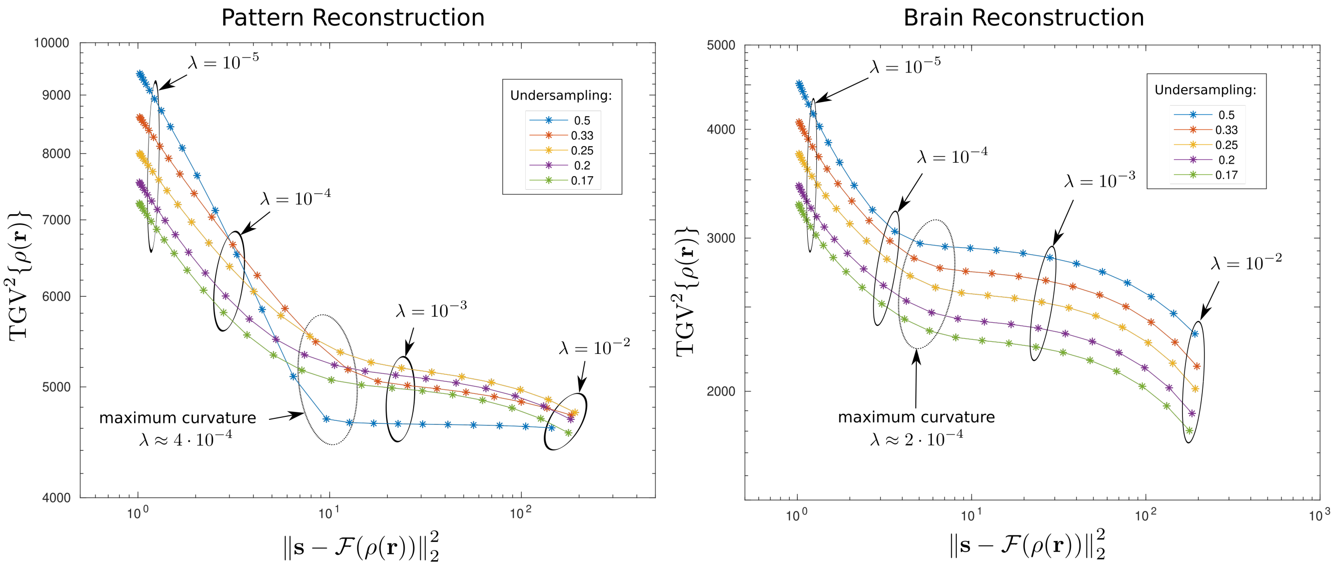


FIG. S6: L-curves of the pattern image and brain image reconstruction. The x-axis represents the energy of the data fidelity term of the solution whereas the y-axis is the TGV regularization energy term. The optimum regularization parameter determined by the NRMSE in fig.S5 is slightly greater than the point of maximal curvature.

### III. ADDITIONAL CONSIDERATIONS ON RECONSTRUCTION PARAMETERS AND RESULTS

The TGV regularization parameter used in the reconstruction was adjusted to $\lambda = 3 \times 10^{-4}$ by gradually increasing the regularization until disappearance of noise-like artifact in the metabolite maps [8]. An L-curve was computed for one of the volunteer dataset ( supplementary Fig.S14) and $\lambda = 3 \times 10^{-4}$ matches the point of maximum curvature that is also considered as the optimal regularization parameter value [1]. The optimal value found was slightly lower than the previously found value in the 2D case ( $\lambda = 10^{-3}$ in [4]). However, this same value was observed to be optimal for all 4 volunteer datasets. Therefore, the regularization parameter seems to depend mainly on the MRSI protocol and might require some minor adjustment when the slab thickness, the resolution, the flip-angle or other acquisition parameters are modified.

The number of components in the model ($K$ in main manuscript eq.(2)) was chosen empirically as follows. The initial estimation of the components were computed with SVD performed on the adjoint solution of the MRSI data. These initial components were then visualized and $K$ was chosen such that all $U_n, V_n$ for $n > K$ contain mainly noise. This upper limit was found to be consistently $\approx 35$ for all subjects and was safely set to 40. For $K < 20$, key metabolite features tend to disappear and there was no noticeable difference in the resulting metabolite maps with $K$ chosen between 40 and 20. Therefore, choosing $K = 40$ was considered safe.

A cautious assessment of the LCModel spectral fit would notice the presence some visible residuals around 3.1 and 3.3 ppm in the fit of the displayed spectra in main manuscript fig.2 and 4. These residuals might be caused by a discrepancy in the peak shape between the reconstructed MRSI datasets and the reference LCModel basis set. A possible cause of this difference might be the 1st order spectral phase correction by estimation of the first FID missing point as described in the methods.

The sample spectra in main manuscript fig.2 and 4 show a particular high Ins peak at 3.9 ppm. Compared to other MRS sequences, ultra-short TE FID-MRSI sequence maximizes the signal of Ins with respect to other metabolites. The Ins signal is the result of overlapping multiplets evolving with strong proton J-coupling. With long TE, the J-coupling results in modulation and dephasing of each multiplet peak, leading to a lower apparent Ins signal at 3.9 ppm. On the contrary, with TE $<$ 1ms in the present study, there is practically no modulation time between the excitation and the acquisition, and the Ins signal is maximum. A simulation illustrating this effect is shown in supplementary Fig.S19.

Cerebellum was only partially covered by the VOI but the results of the anatomical analysis are in agreement with previously reported data that showed a high levels of tCre, Glx and Ins in this structure of the hindbrain [9, 10]. With the presented acquisition over the whole-brain, an extension of the VOI to cover the entire cerebellum would cause a degradation of the data quality because of shimming performance limitations. Nevertheless, recent work showed the feasibility of acquiring MRSI over the whole cerebellum [11]

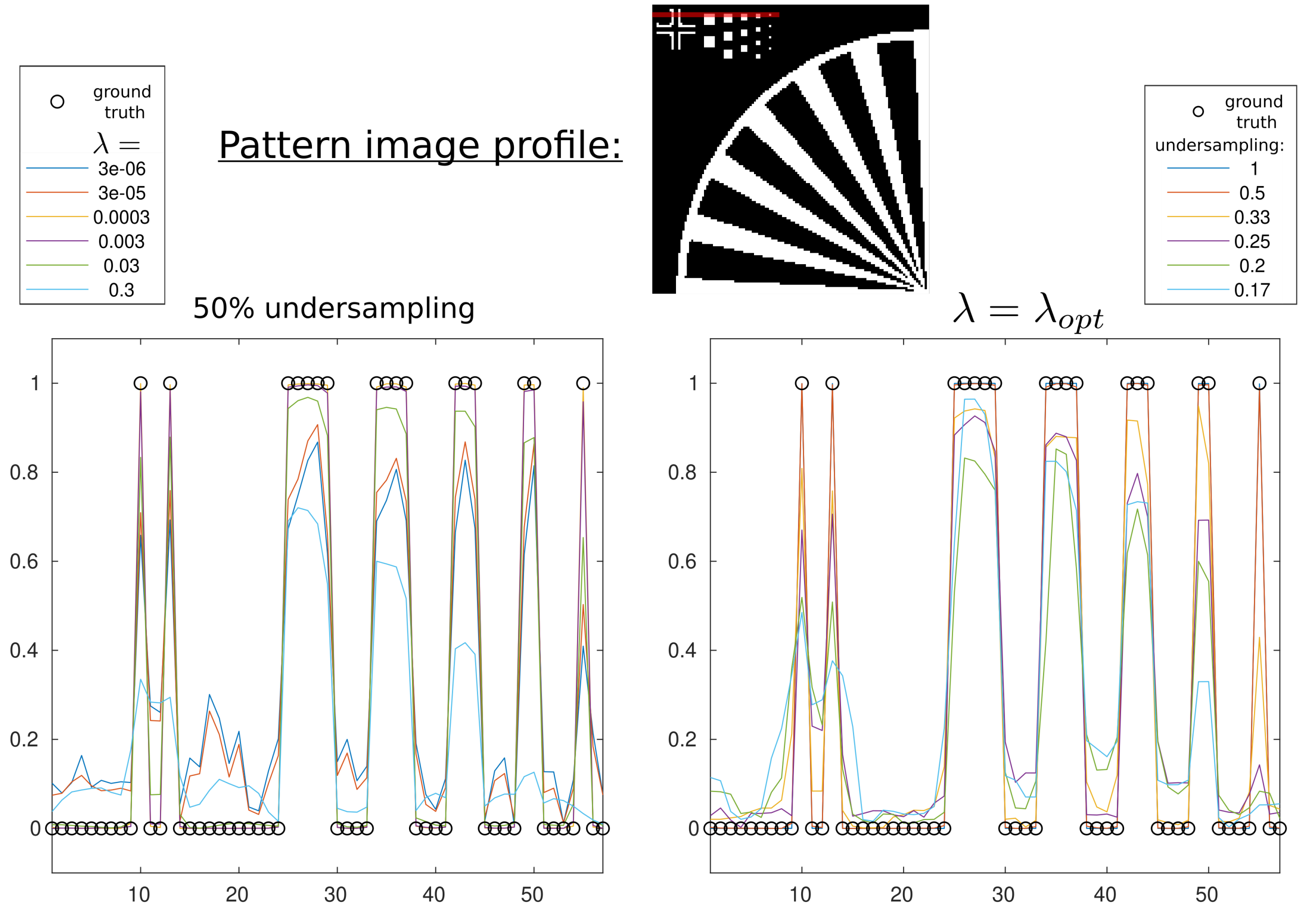


FIG. S7: Analysis of a profile in the pattern image (profile location shown with the red line on the top image). Left, reconstruction results are shown for 50% k-space undersampling and varying TGV regularization parameter, $\lambda$. Right, the regularization parameter is set to its optimal value $\lambda_{opt} = 1.5 \cdot 10^{-3}$ but with a varying k-space undersamping factor. The reference image value of the profile are shown with the black circles.

### IV. SUPPLEMENTARY FIGURES


[1] H. W. Engl and W. Grever, Numerische Mathematik **69**, 25 (1994), ISSN 0029-599X, URL `http://link.springer.com/10.1007/s002110050078`.

[2] D. Donoho, IEEE Transactions on Information Theory **52**, 1289 (2006), ISSN 0018-9448.

[3] L. Michael, D. David, and P. J. M., Magnetic Resonance in Medicine **58**, 1182 (2007).

[4] A. Klauser, S. Courvoisier, J. Kasten, M. Kocher, M. Guerquin-Kern, D. Van De Ville, and F. Lazeyras, Magnetic Resonance in Medicine **81**, 2841 (2019), ISSN 07403194.

[5] J. A. Maldjian, P. J. Laurienti, R. A. Kraft, and J. H. Burdette, NeuroImage (2003), ISSN 10538119.

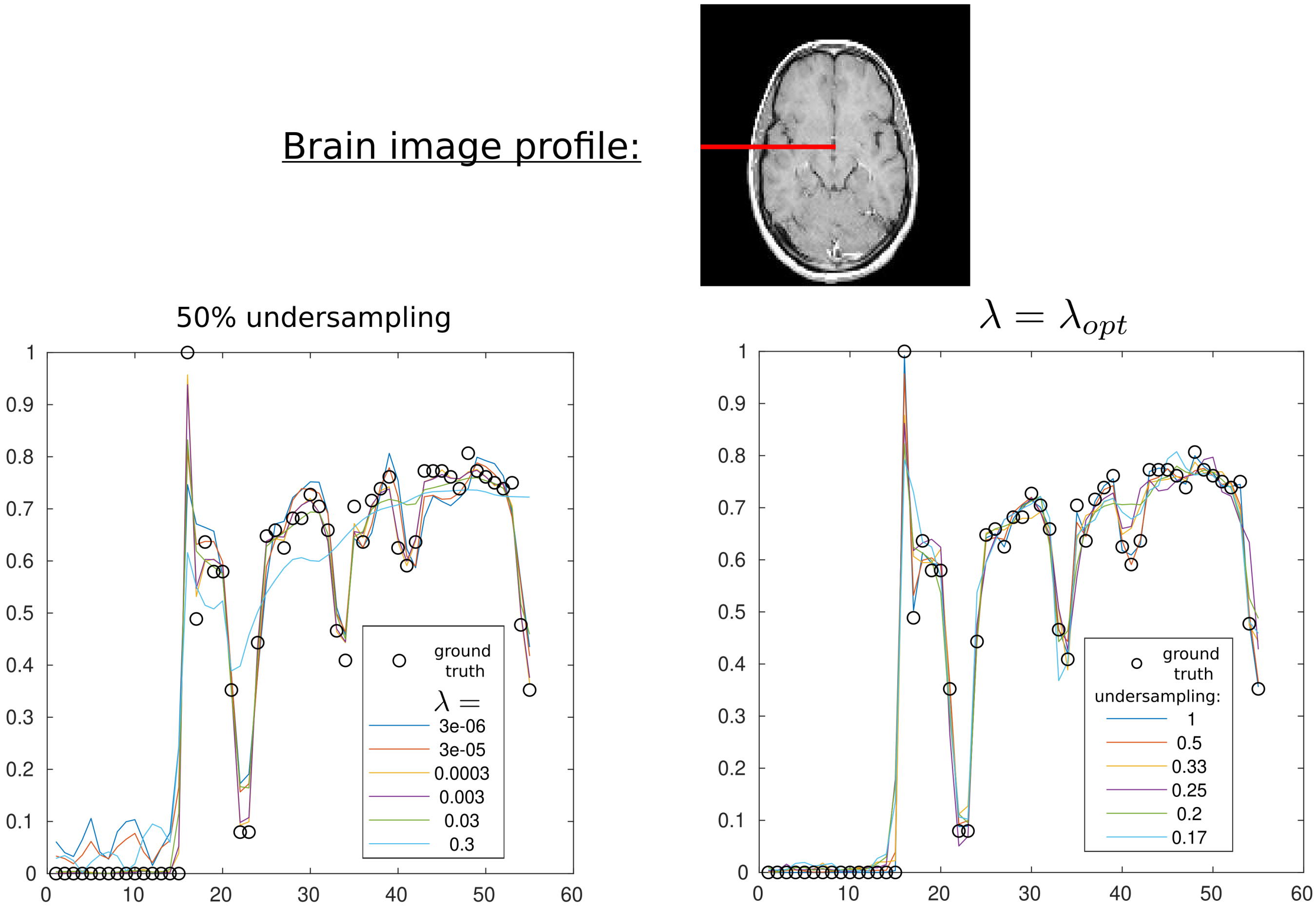


FIG. S8: Analysis of a profile in the brain image (profile location shown with the red line on the top image).Left, reconstruction results are shown for 50% k-space undersampling and varying TGV regularization parameter, $\lambda$. Right, the regularization parameter is set to its optimal value $\lambda_{opt} = 5 \cdot 10^{-4}$ but with a varying k-space undersamping factor. The reference image value of the profile are shown with the black circles

[11] U. E. Emir, J. Sood, M. Chiew, M. A. Thomas, and S. P. Lane, Magnetic Resonance in Medicine **85**, 2349 (2021), ISSN 0740-3194, URL `https://onlinelibrary.wiley.com/doi/10.1002/mrm.28614`.

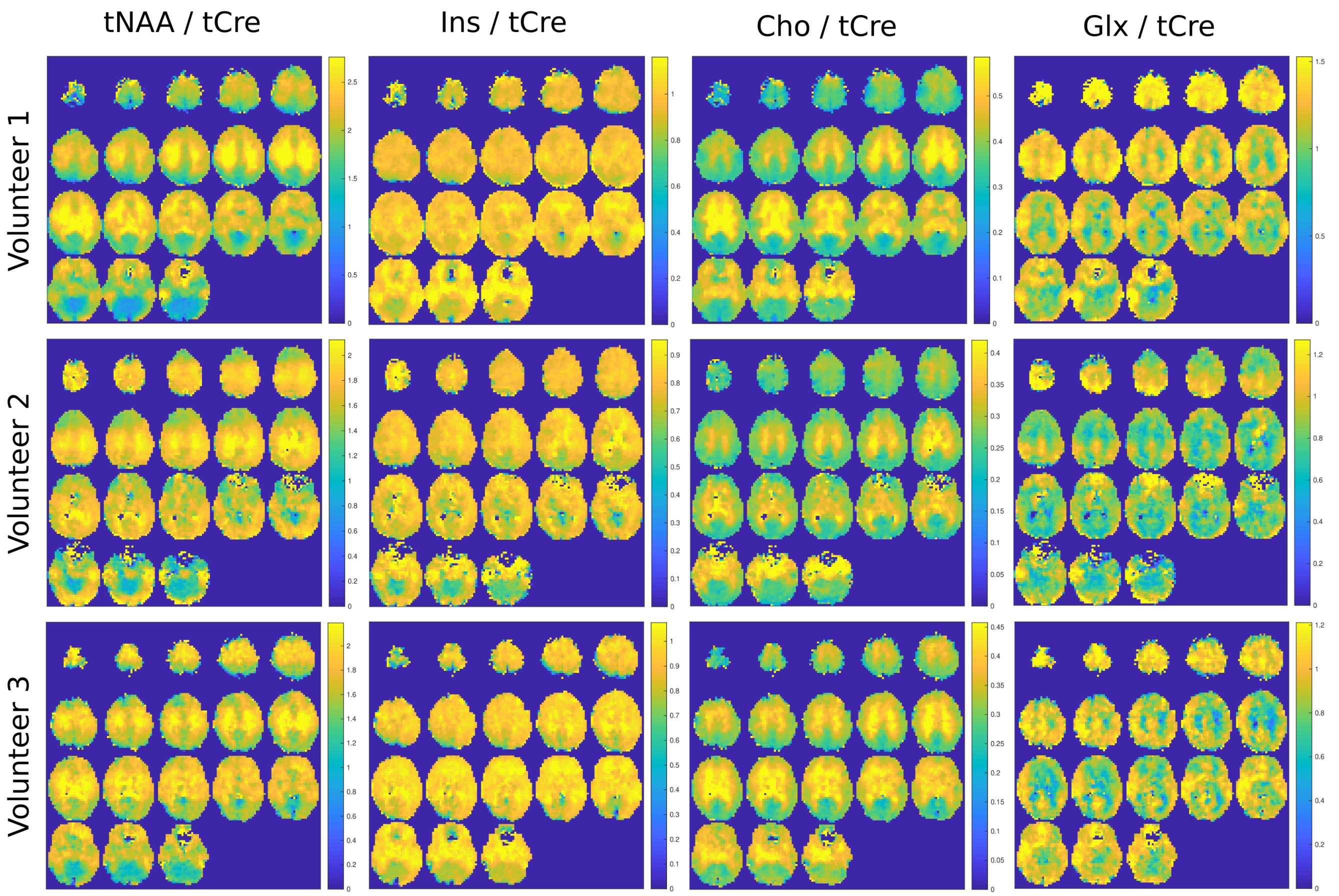


FIG. S9: Metabolite concentration ratio maps with respect to tCre. The individual metabolite maps are presented in main manuscript fig.4 for the three volunteers.

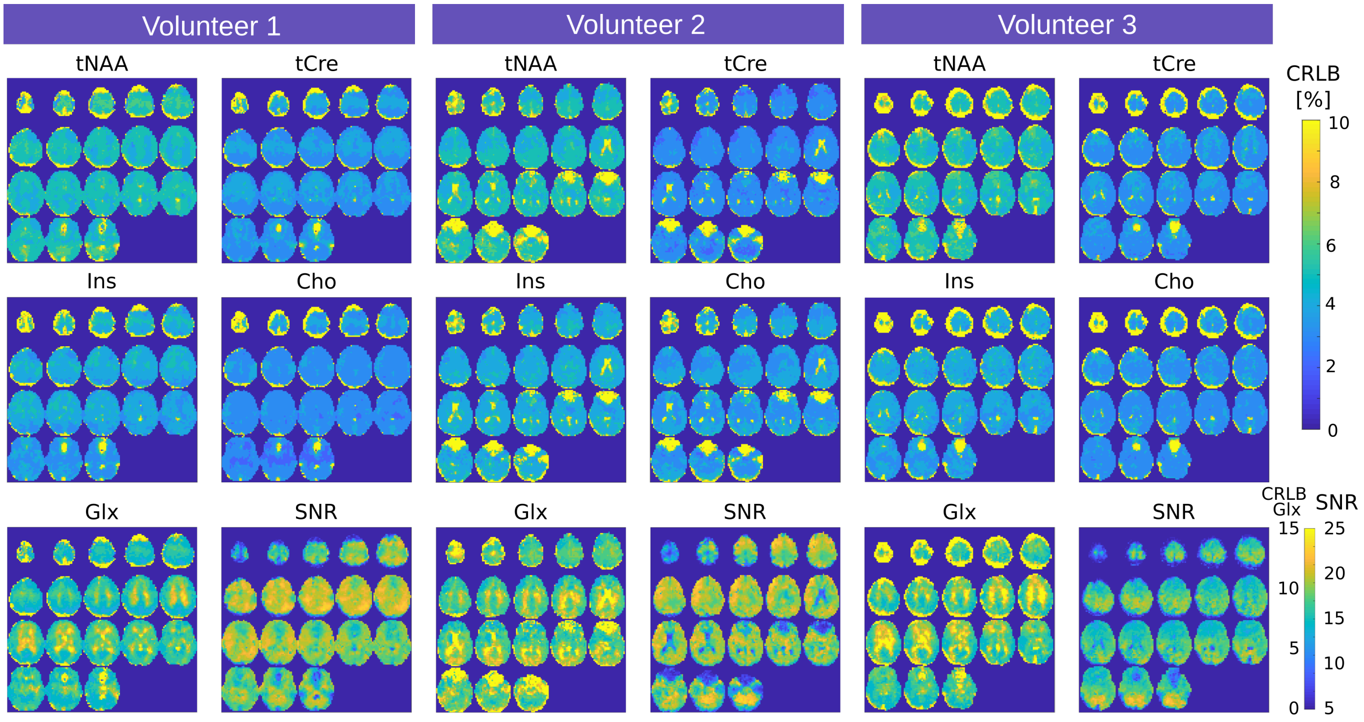


FIG. S10: Cramer-Rao lower bound (CRLB) and signal-to-noise ratio (SNR) as estimated by LCModel for the metabolite maps of the three volunteers presented in main manuscript fig.4. CRLB of tNAA, tCre, Ins, Cho is presented with a colorbar ranging from 0 to 10%. Colorbar of Glx CRLB ranges from 0 to 15%. The SNR maps are shown with a range from 0 to 25.

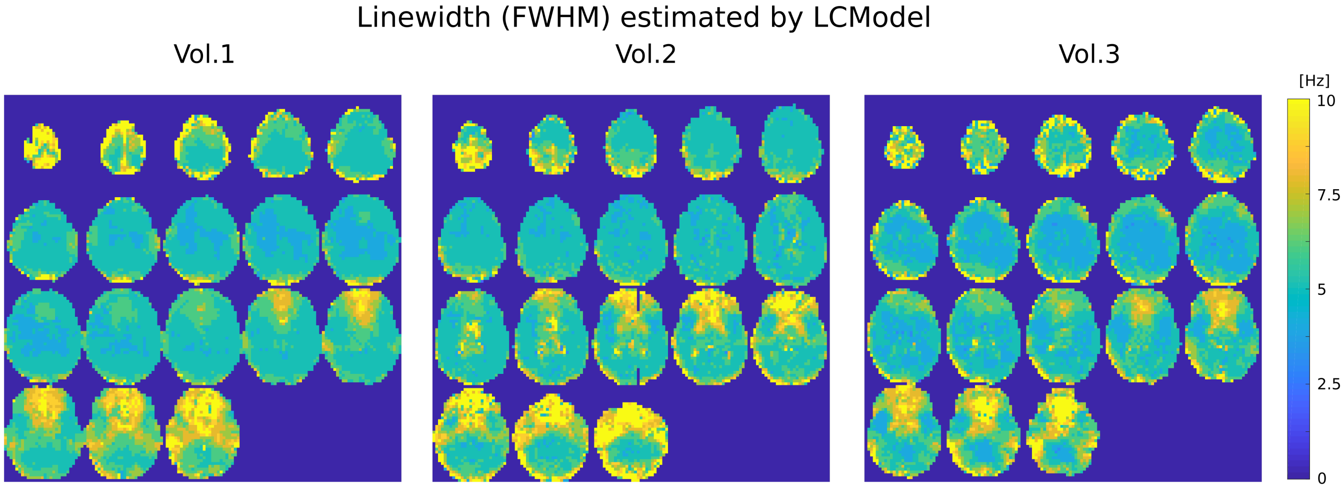


FIG. S11: Linewidth maps of the three volunteers 3D MRSI reconstructions presented in main manuscript fig.4.

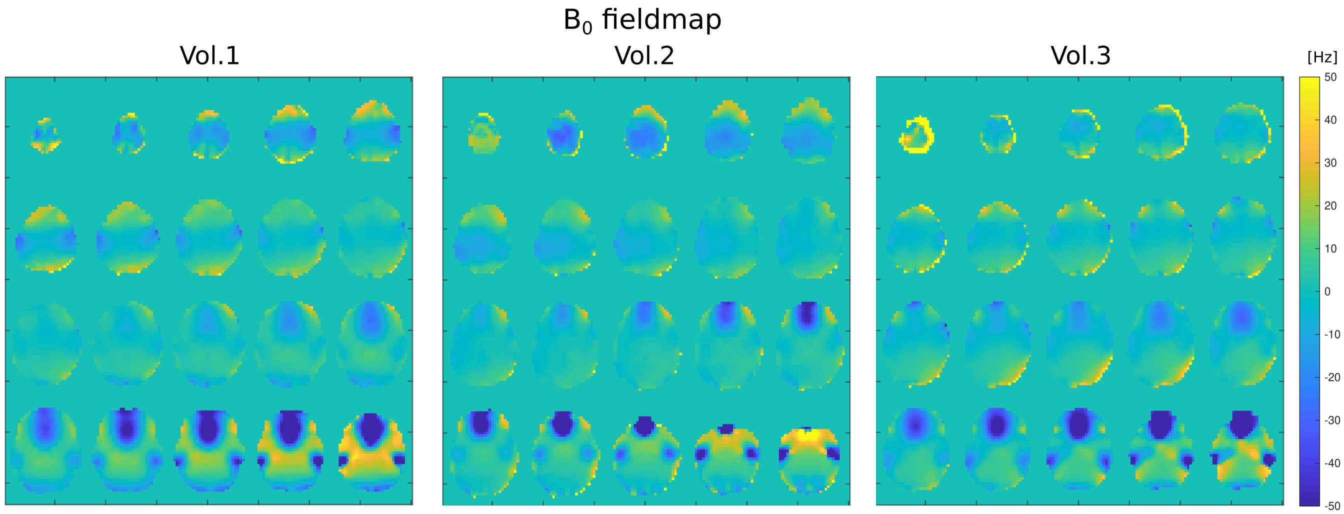


FIG. S12: $B_0$ fieldmap measured from water reference data for the three volunteers 3D MRSI acquisitions presented in main manuscript fig.4.

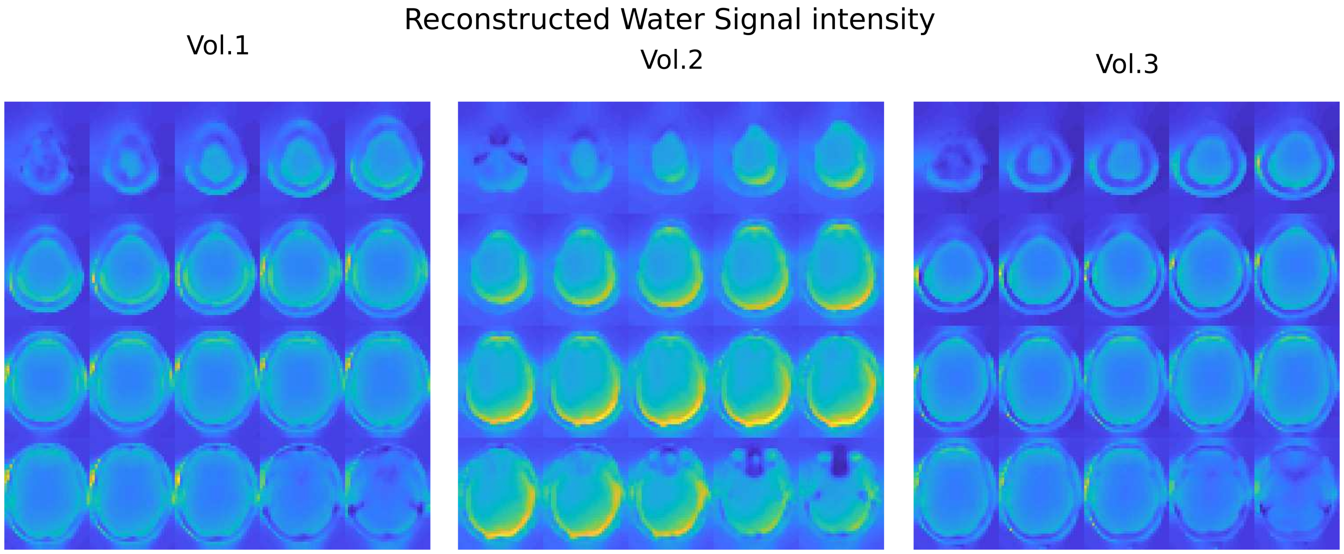


FIG. S13: Water signal reconstructed for reference corresponding to the three volunteers 3D MRSI acquisitions presented in main manuscript fig.4.

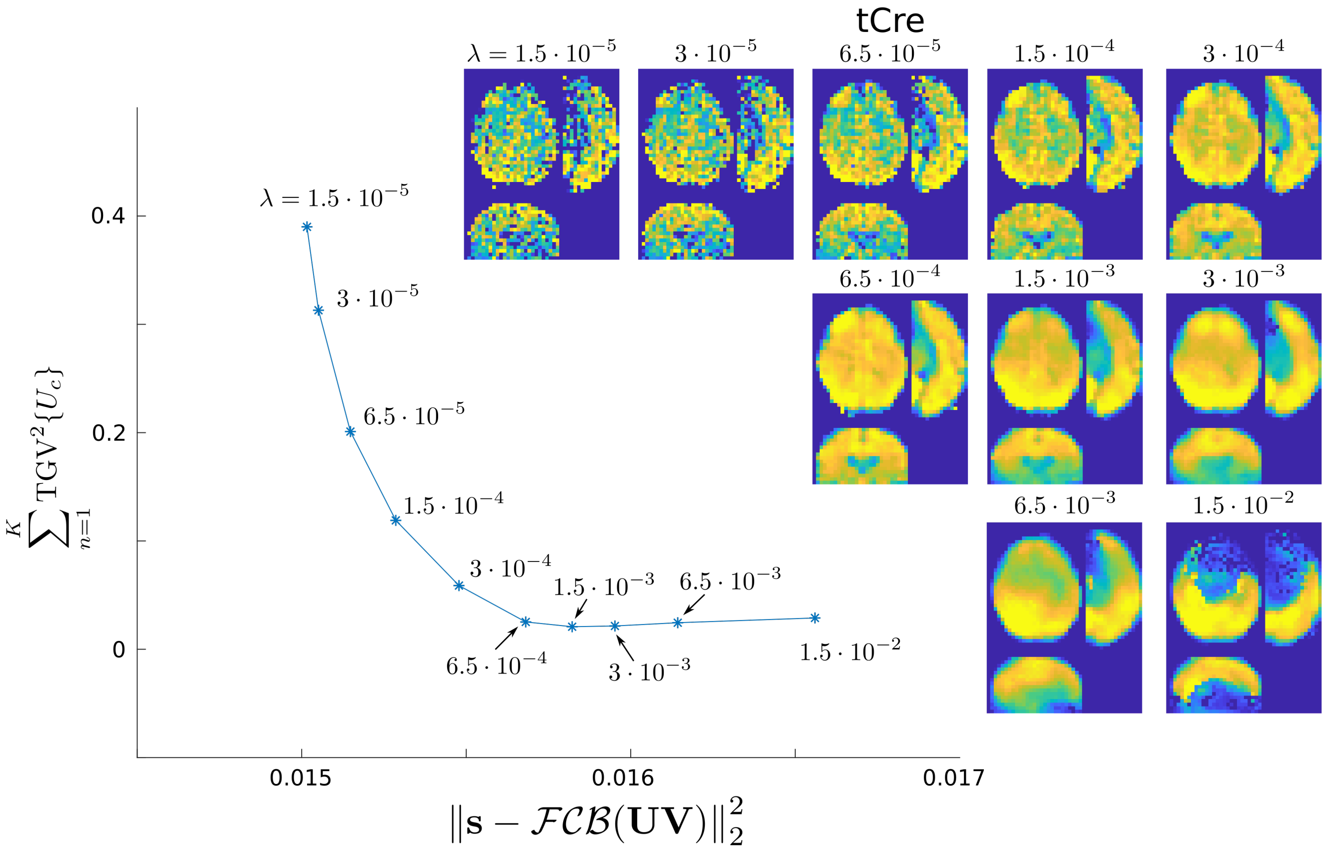


FIG. S14: L-curve performed with healthy volunteer MRSI Data. The regularization parameter value used for the main manuscript data ($\lambda = 3 \cdot 10^{-4}$) corresponds to the point of maximum curvature that is considered as the optimal regularization parameter value.

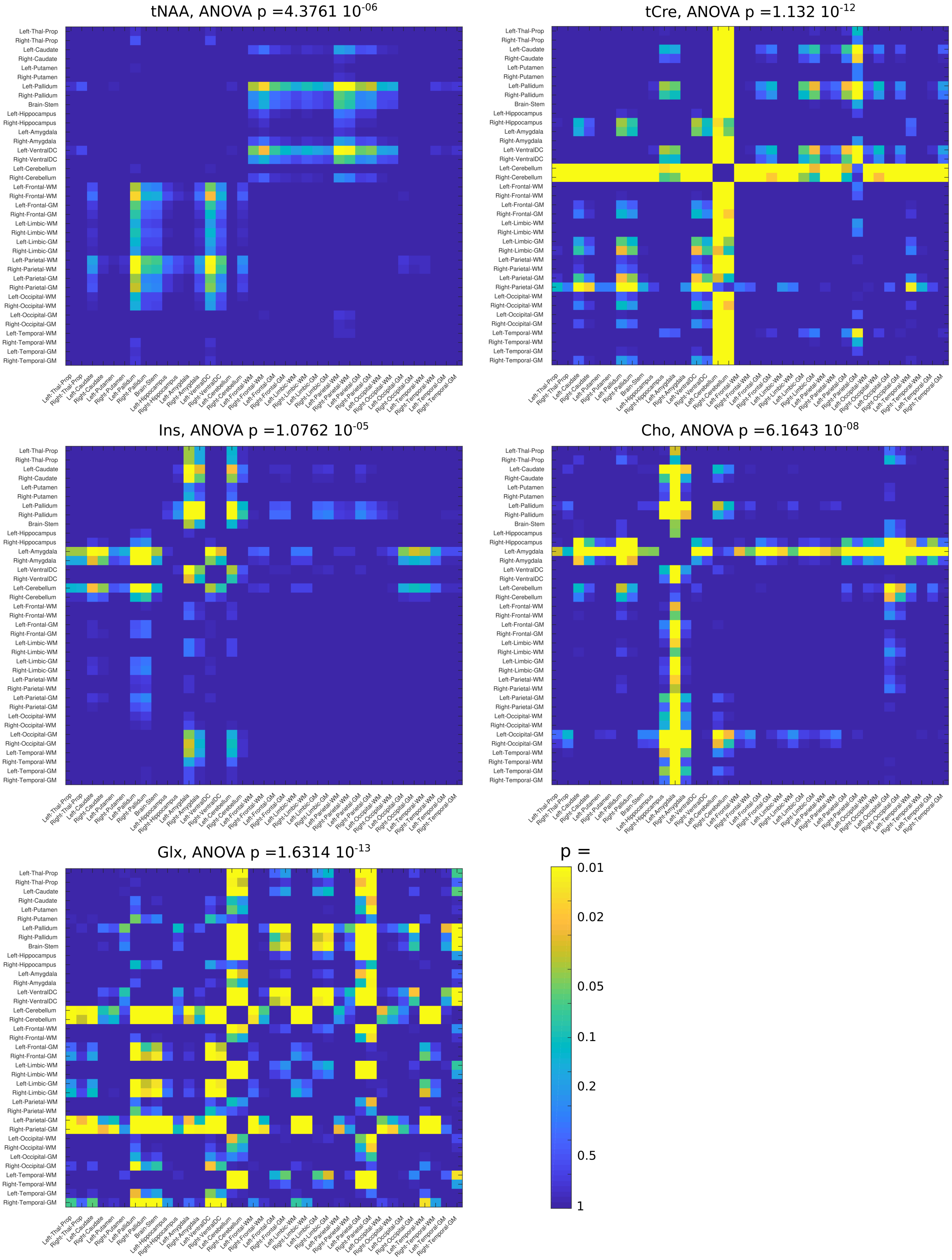


FIG. S15: Statistical testing of the anatomical segmentation results presented in the main manuscript (fig. 6). For each metabolite, the title indicates the p-value from the ANOVA testing that was significant for all metabolite. Multiple comparison across region was performed using Tukey's test and results are shown on matrices with the brightest pointing the most significant difference.

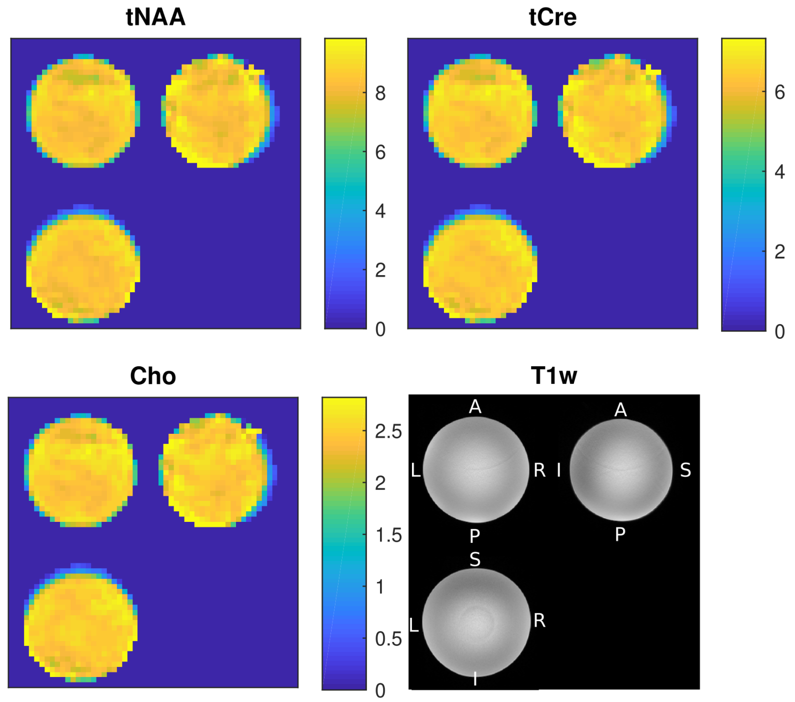


FIG. S16: 3D FID-MRSI measurement with CS Acceleration factor = 3.5 and 5 mm isotropic resolution on a spherical phantom with homogenous metabolite solution. The CS-SENSE-LR resulting metabolite maps (in institutional units) aim to demonstrate the ability of the method to reconstruct an homogenous concentration distribution with the correct water reference signal.

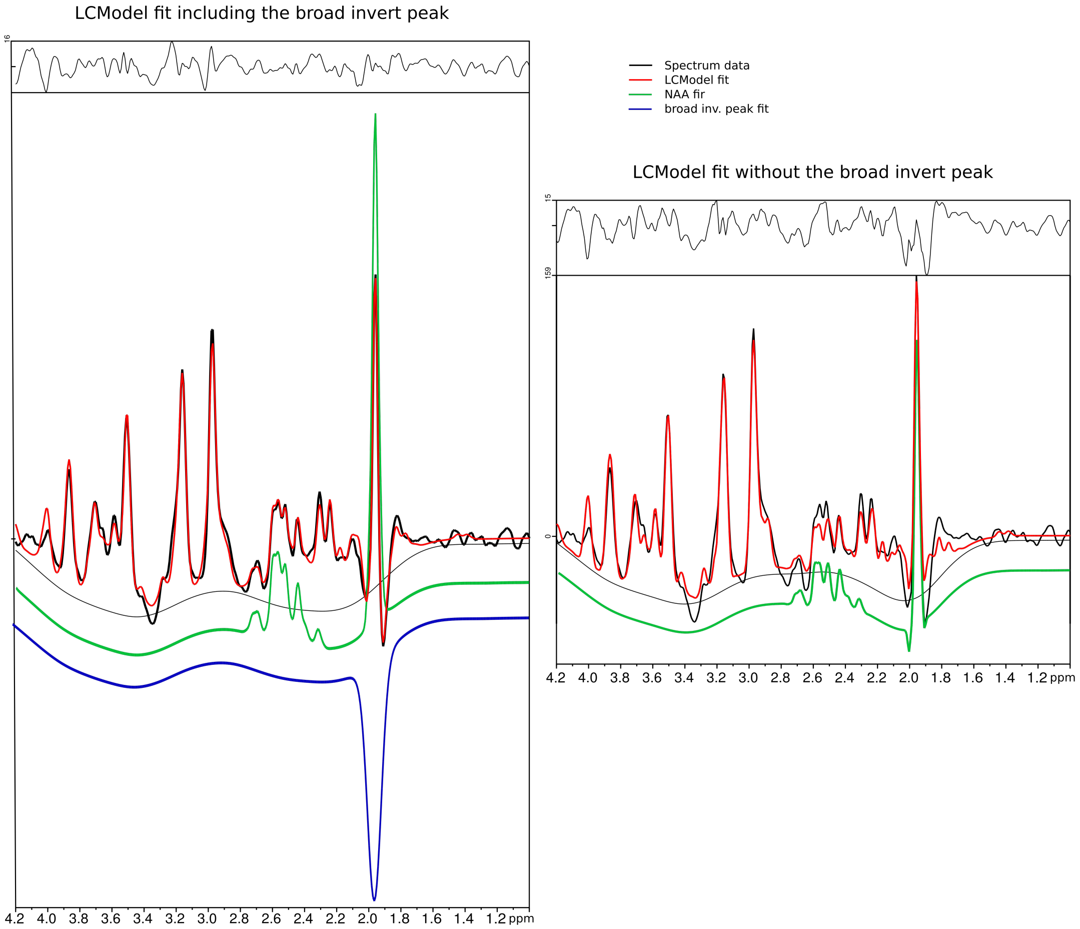


FIG. S17: LCModel fit of a sample spectrum with or without the 20Hz-broad invert peak at 2 ppm. The fit residuals are clearly reduced at 2 ppm in presence of the invert peak in the fit. Without the invert peak, NAA is clearly underestimated as observable with the multiplet at 2.5 ppm, far from the lineshape distortion. Glx at 2.3 ppm seem also underestimated in absence of invert peak.

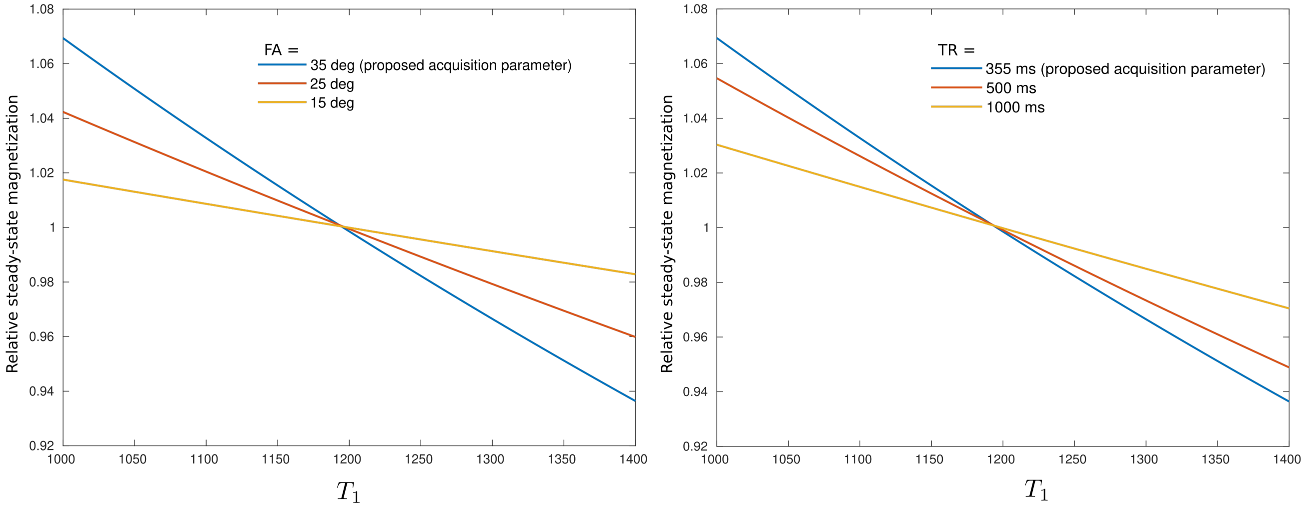


FIG. S18: Calculation of the steady-state magnetization as function of the $T_1$ relaxation time for several flip-angles (FA) and repetition times (TR). The transverse steady-state magnetization is given by $M_{steady}(T_1) = M_0^z \frac{\sin(FA)\left(1-e^{-TR/T_1}\right)}{1-\cos(FA)e^{-TR/T_1}}$, with $M_0^z$ the longitudinal resting-state magnetization. The plots represent the relative magnetization $\frac{M_{steady}(T_1)}{M_{steady}(1200ms)}$

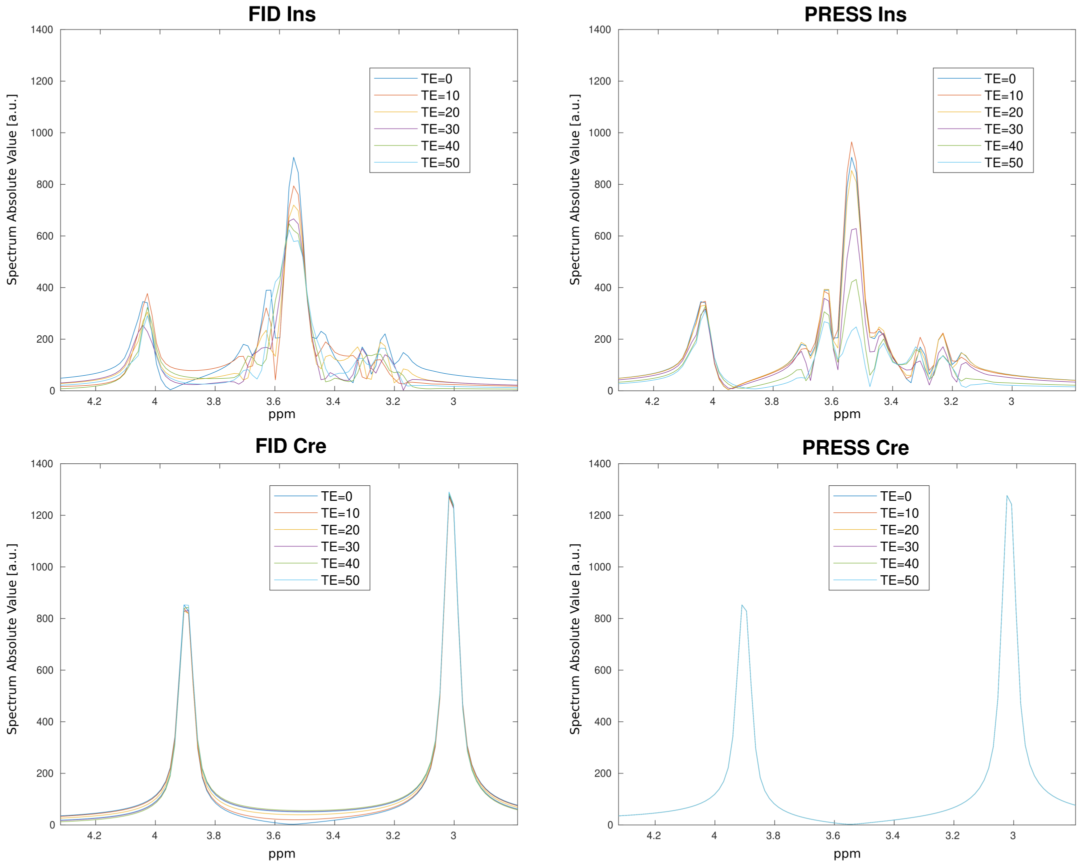


FIG. S19: Simulation of myo-inositol (Ins) and creatine (Cre) signal for FID (excite-acquire) and PRESS sequence for different TE. The PRESS sequence was simulated with ideal pulses (perfect refocusing pulses with no duration) and T2 relaxation was not simulated. The goal here is to show the effect of Ins multiplet J-coupling on the resulting signal amplitude. The major Ins peak is a contribution of 2 overlapping doublets. The apparent loss of Ins signal with longer TE in both sequences is due to the J-coupling modulation of the doublets occuring with time. The longer the TE, the more dephased are the multiplets components overlapping at 3.55 ppm and the lower is the apparent resulting Ins peaks at 3.55 ppm. For comparison, Cre peaks at 3.9 and 3 ppm are actual singlets. Therefore, their amplitudes are not affected by the TE (ommiting the T2 effect for sake of the demonstration) neither for FID or PRESS sequence.

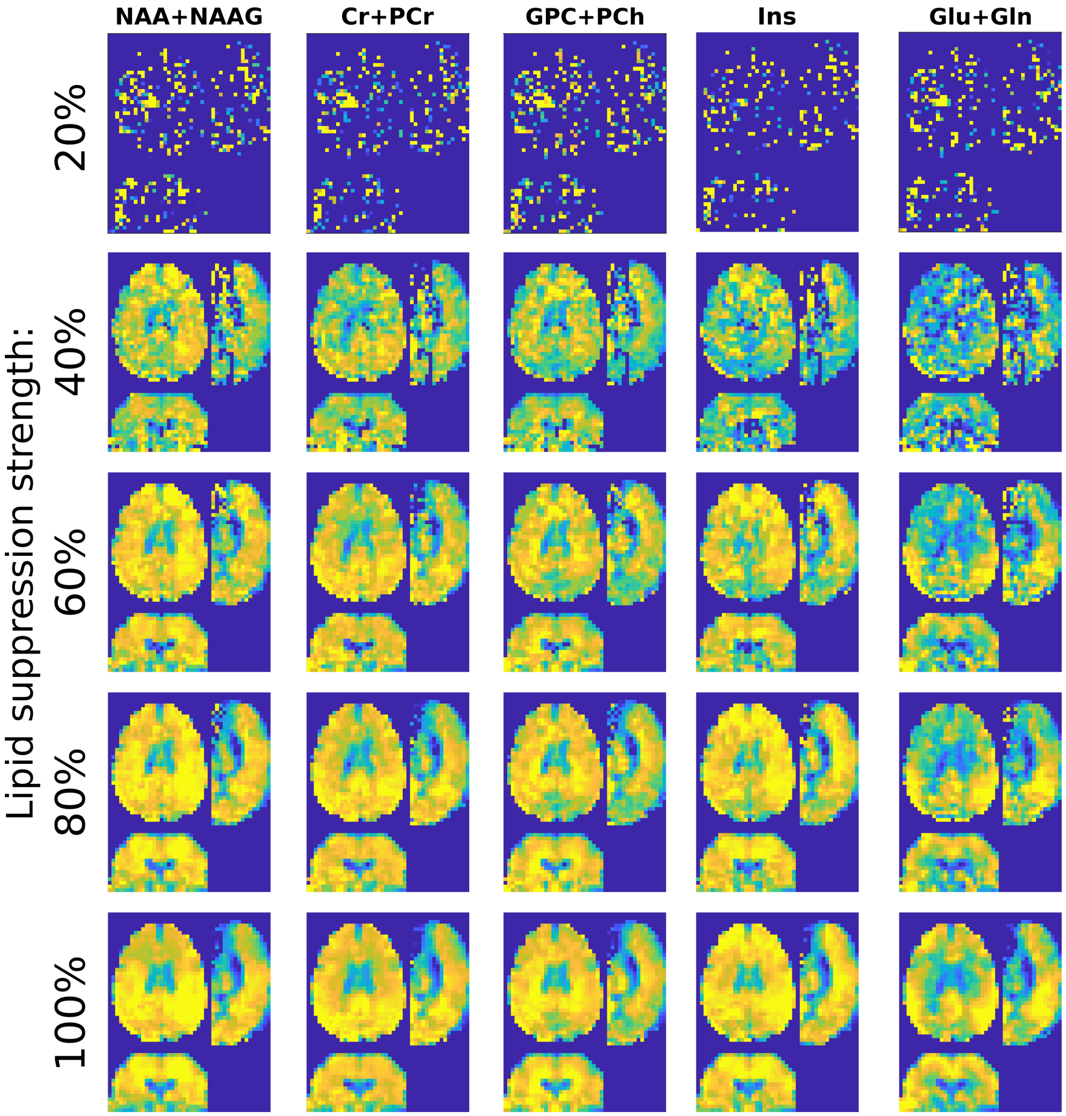


FIG. S20: Effect of reducing the strength of the lipid suppression by orthogonality. The strength was reduced by gradually reducing the criteria for determination of the lipid suppresion rank [4]. The contamination of the resulting metabolite maps by the lipid is visible for all metabolites and for any strength $< 100\%$.
